# Visualizing Mitochondrial Heme Flow through GAPDH to Targets in Living Cells and its Regulation by NO

**DOI:** 10.1101/2024.01.10.575067

**Authors:** Pranjal Biswas, Joseph Palazzo, Simon Schlanger, Dhanya Thamaraparambil Jayaram, Sidra Islam, Richard C. Page, Dennis J. Stuehr

## Abstract

Iron protoporphyrin IX (heme) is an essential cofactor that is chaperoned in mammalian cells by GAPDH in a process regulated by NO. To gain further understanding we generated a tetra-Cys human GAPDH reporter construct (TC-hGAPDH) which after being expressed and labeled with fluorescent FlAsH reagent could indicate heme binding by fluorescence quenching. When purified or expressed in HEK293T mammalian cells, FlAsH-labeled TC-hGAPDH displayed physical, catalytic, and heme binding properties like native GAPDH and its heme binding (2 mol per tetramer) quenched its fluorescence by 45-65%. In live HEK293T cells we could visualize TC-hGAPDH binding mitochondrially-generated heme and releasing it to the hemeprotein target IDO1 by monitoring cell fluorescence in real time. In cells with active mitochondrial heme synthesis, a low-level NO exposure increased heme allocation into IDO1 while keeping steady the level of heme-bound TC-hGAPDH. When mitochondrial heme synthesis was blocked at the time of NO exposure, low NO caused cells to reallocate existing heme from TC-hGAPDH to IDO1 by a mechanism requiring IDO1 be present and able to bind heme. Higher NO exposure had an opposite effect and caused cells to reallocate existing heme from IDO1 to TC-hGAPDH. Thus, with TC-hGAPDH we could follow mitochondrial heme as it travelled onto and through GAPDH to a downstream target (IDO1) in living cells, and to learn that NO acted at or downstream from the GAPDH heme complex to promote a heme reallocation in either direction depending on the level of NO exposure.

## Introduction

In eukaryotes iron protoporphyrin IX (heme) is generated within the mitochondria and must be exported and transported from this organelle to reach the numerous proteins that require heme to mature and function in other cellular compartments. Due to the physical and chemical properties of heme, its transport in cells has long been expected to involve a protein chaperone (1-5). Recently, the glycolytic enzyme GAPDH was shown to fulfill this role by binding mitochondrial heme and enabling its delivery to diverse targets including hemoglobin α, β, and γ, myoglobin, tryptophan dioxygenase (TDO), indoleamine dioxygenase 1 (IDO1), soluble guanylyl cyclase (sGC), NO synthases, and heme oxygenase 2 (6-12). In most cases, the heme insertions into these proteins also required the cell chaperone Hsp90 and its ATPase activity (13). Thus, GAPDH and Hsp90 have emerged as important players for intracellular heme allocation to diverse targets in mammalian cells.

Cells use multiple systems to carefully regulate their heme levels (14), but how their heme levels relate to internal heme distribution is mostly unclear. Evidence suggests that outside of the erythroid system, cell heme availability is naturally kept in deficit and this results in cell heme proteins being only partially heme-saturated under normal culture conditions (15). Within this context, the signaling molecule NO has emerged as an important regulator of intracellular heme allocation (15,16). Specifically, relatively low level NO exposure promoted cell heme allocation and thus caused the heme content of several heme proteins to increase by a process that also depended on GAPDH and Hsp90. In contrast, at higher NO exposure levels the promoting effect of NO was quickly lost and instead it widely prevented cellular heme allocation to proteins (17,18) and even caused some to lose their heme (17). NO displayed this hormetic effect when it was released from small molecule NO donors or when it was naturally generated in cells via NO synthases (17,19-21). Overall, this suggests a new way that NO can shape biological processes by positively or negatively regulating cell heme allocation and the consequent functions of numerous heme-dependent proteins.

Thus far, our understanding of cell heme allocation has been limited by an inability to follow mitochondrial heme as it travels onto GAPDH and from it to its downstream targets. We therefore sought a means to monitor the GAPDH heme binding and transfer events in living cells and to study how they may be regulated by NO. We designed a reporter construct called tetra-Cys human GAPDH (TC-hGAPDH) which, following its expression and labeling with a commercially-available FlAsH reagent (22), we envisioned could indicate GAPDH heme binding by fluorescence quenching when in its purified form and when it is expressed in mammalian cells. Here, we characterize TC-hGAPDH, demonstrate its utility for visualizing GAPDH heme binding in living cells and heme transfer from GAPDH to the client protein IDO1, and for investigating the impact of NO on these heme transfer steps. We expressed IDO1 and TC-hGAPDH in HEK293T cells under two conditions: When the cells were cultured normally and thus contained a normal level of heme, and when the cells were cultured with heme precursors δ-ALA and ferric citrate (Fe-cit) to stimulate mitochondrial heme synthesis and generate a supranormal level of heme in the cells (8,23). In addition, we investigated how turning off cell mitochondrial heme biosynthesis at the point of initiating low or high NO exposures would impact the NO effect on heme distributions in TC-hGAPDH and IDO1. Our findings provide a first look at how mitochondrially-generated heme flows through GAPDH to a target heme protein (IDO1) in living cells, how NO regulates their heme distributions, and reveal that a NO-driven heme exchange can occur between the two proteins in either direction depending on the level of NO exposure. Overall, this demonstrates how TC-hGAPDH can help to better understand cell allocation of mitochondrial heme and its regulation by NO.

## Results

### Generation and characterization of TC-hGAPDH

To enable monitoring of GAPDH heme binding we adopted FIAsH-based technology (22) like we successfully utilized before to study heme binding by the hemeprotein sGCβ (19). This involved inserting a short DNA sequence into our human GAPDH expression plasmids that encode for a FlAsH biarsenical-binding tetracysteine (TC) motif immediately following His57 (Fig. 1A). This location is in a loop region downstream from the presumed heme-coordinating His53 in GAPDH (24), thus creating TC-hGAPDH (sequence details in Fig. S1). The TC motif is designed to bind the bi-arsenical dye FIAsH (Fig. 1B), which only fluoresces after it becomes bound to the TC sequence in a protein (22,25). The TC-hGAPDH construct we designed for bacterial expression also contained a C-terminal GST fusion protein to aid in purification (24). The TC-hGAPDH construct we designed for mammalian cell expression contained an N-terminal HA tag as we utilized before (7) to facilitate its IP pull-down and to allow us to distinguish it from endogenous GAPDH present in cells.

**Figure 1.**
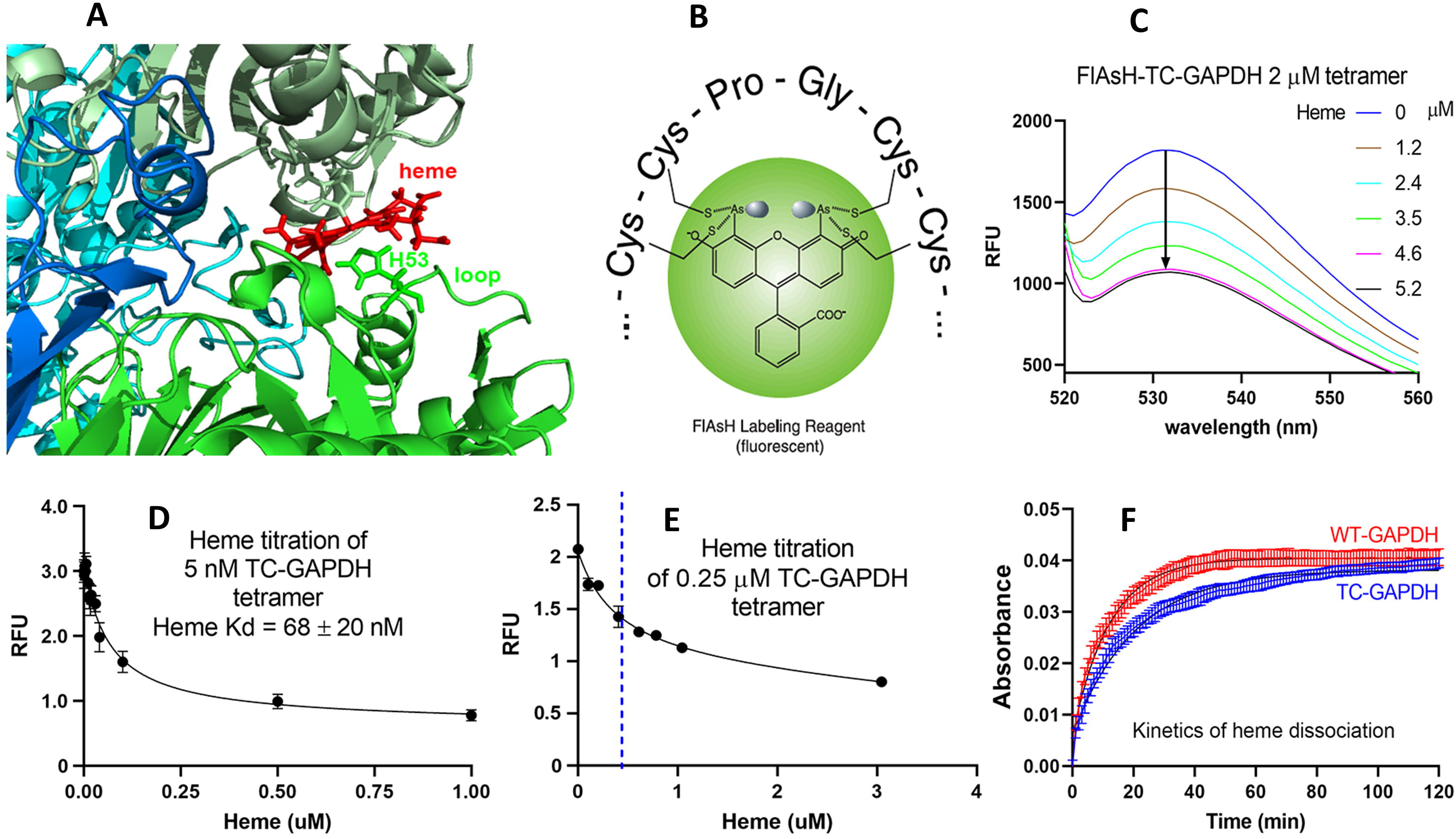
TC-hGAPDH and its heme binding properties. *Panel A-* Close-up of a representative structure from molecular dynamics simulations of the modeled human GAPDH–heme complex, with the loop for TC insertion indicated. Adapted from Sweeny et. al., JBC 2018. *Panel B-* Structure of the FlAsH reagent bound to a prototypical TC amino acid sequence. *Panel C-* Fluorescence quenching of bacterially expressed, purified, FlAsH-labeled TC-hGAPDH upon titration with ferric heme. *Panel D-* Representative ferric heme titration of FlAsH-labeled TC-hGAPDH to determine the Kd. *Panel E-* Representative ferric heme titration of FlAsH-labeled TC-hGAPDH, with the dashed blue line indicating two heme added per tetramer. *Panel F-* Representative graph comparing the kinetics of ferric heme dissociation from purified wild type hGAPDH or TC-hGAPDH upon mixing them with excess apo-myoglobin.

Our analysis of the bacterially-expressed and purified TC-hGAPDH indicated the TC insertion did not destabilize hGAPDH tetrameric structure as determined by analytical gel filtration chromatography (data not shown) nor did it change the enzyme dehydrogenase activity (25 ± 3 *versus* 26 ± 1 mU/ml for wild type and TC-hGAPDH, respectively, n = 3). Titrating FIAsH-labeled TC-hGAPDH with ferric hemin quenched its fluorescence by 44 ± 7% (n = 3) at apparent heme saturation (Fig. 1C). Equilibrium titrations with ferric hemin gave a Kd of 68 ± 20 nM for TC-hGAPDH (Fig. 1D) and indicated that it contained two high-affinity heme binding sites per tetramer (Fig. 1E), consistent with our structural model of the GAPDH-heme complex with two identical heme sites per tetramer (24). Heme dissociation experiments in the presence of excess apo-myoglobin were done at room temperature and gave a rate of ferric heme dissociation (k_off_) from FlAsH-TC-hGAPDH of 9.1 ± 1.4 x 10^-4^ s^-1^, n = 3 (Fig. 1F), which was similar to the heme k_off_ we measured for human GAPDH under the same conditions (9.7 ± 1.0 x 10^-4^ s^-1^, n = 3). Thus, incorporating the TC insertion into hGAPDH did not significantly alter its heme binding, catalytic, or physical properties, thus suggesting that TC-hGAPDH could be a useful probe to indicate and follow GAPDH heme binding in cells.

### HA-TC-hGAPDH heme binding in mammalian cells

HEK293T cells that had been made heme deficient by culturing for 2 d with succinyl acetone (SA) and in heme-depleted serum (24) were transfected to express the HA-TC-hGAPDH protein. Expression under this culture condition ensured that the HA-TC-hGAPDH would be as heme-free as possible. Following FlAsH labeling and cell lysis, the cell supernatant was titrated with hemin and gave a Kd of 100 ± 25 nM, n = 6 for the HA-TC-hGAPDH protein and resulted in 65 ± 2% (n = 5) quenching of its fluorescence at saturation (Fig. S2), which is similar to the Kd and percent quenching values that we observed for the bacterially-expressed and purified TC-hGAPDH as reported above. Thus, HA-TC-hGAPDH expressed well in mammalian cells and upon FlAsH tagging displayed heme binding and fluorescence quenching properties consistent with the bacterially-expressed and purified form.

We next investigated how the extent of heme binding by FlAsH-tagged HA-TC-hGAPDH expressed in the heme-deficient HEK293T cells would be impacted when the cells underwent further culture in one of four different conditions of heme availability: Continued full heme deficiency (media containing SA and 10% heme-depleted serum); partial heme deficiency (Medium containing SA and 10% normal serum, which typically results in heme being present at a final concentration of 0.2-0.5 µM due to the serum); normal culture conditions (medium without SA and with 10% normal serum, a condition that also allowed the cells to recover their mitochondrial heme biosynthesis from SA inhibition); and conditions of increased intracellular heme production (medium without SA plus normal serum and containing the mitochondrial heme biosynthesis precursors δ-ALA and Fe-cit) (6,8,23). We then monitored the FlAsH fluorescence signals of HA-TC-hGAPDH in real time in the live cells under each condition in a 96-well plate reader, where T = 0 indicates the point where the cell culture media was exchanged to create the four different conditions of heme availability as described above.

As shown in Fig. 2A, the initial fluorescence intensities at T = 0 were approximately the same for all four cell groups, as expected and consistent with the HA-TC-hGAPDH protein being expressed at a similar level, which we confirmed by Western analysis (Fig. S3). As time progressed the fluorescence intensity remained stable only in the cell group that remained in a fully heme-deficient condition, whereas the fluorescence signals gradually diminished for the other three groups over a 100-150 min period until reaching new equilibrium levels. The kinetics of the HA-TC-hGAPDH fluorescent signal change in the cells that were given δ-ALA and Fe-cit correlated with their undergoing a general increase in intracellular heme level as judged in companion experiments with HEK293T cells that were transfected to express a cytosolic heme sensor protein HS1 (26) in place of HA-TC-hGAPDH (Fig. 2B and S4). When the experiment was repeated using cells that expressed the HA-TC-hGAPDH H53A variant, which has a 30-fold poorer heme binding affinity (24), the addition of δ-ALA and Fe-cit to the cells did not elicit any decrease in FlAsH fluorescence (Fig. 2C), confirming that the fluorescence decreases we observed for cells expressing HA-TC-hGAPDH were due to it binding intracellular heme. The relative extents of heme binding by HA-TC-hGAPDH under the four different conditions of heme availability, as indicated by the fluorescence emission levels after equilibrium was reached in each case in Fig. 2A, had a rank order of δ-ALA/Fe-cit supplemented >> normally cultured cells > normally cultured cells + heme deficient serum > fully heme-deficient cells. This rank order correlated directly with the different levels of total porphyrin or total heme that were in the cell supernatants at the end of each culture condition (Fig. 2D) and with the amounts of ^14^C-heme that were bound to HA-TC-hGAPDH or its H53A variant as determined by IP pulldowns of the HA-tagged proteins from replica experiments that utilized ^14^C-δ-ALA (Fig. 2E). These results confirm that the FlAsH fluorescence signal is a good indicator of heme binding by HA-TC-hGAPDH in live cells and can be used to visualize the kinetics and extent of its heme binding.

**Figure 2.**
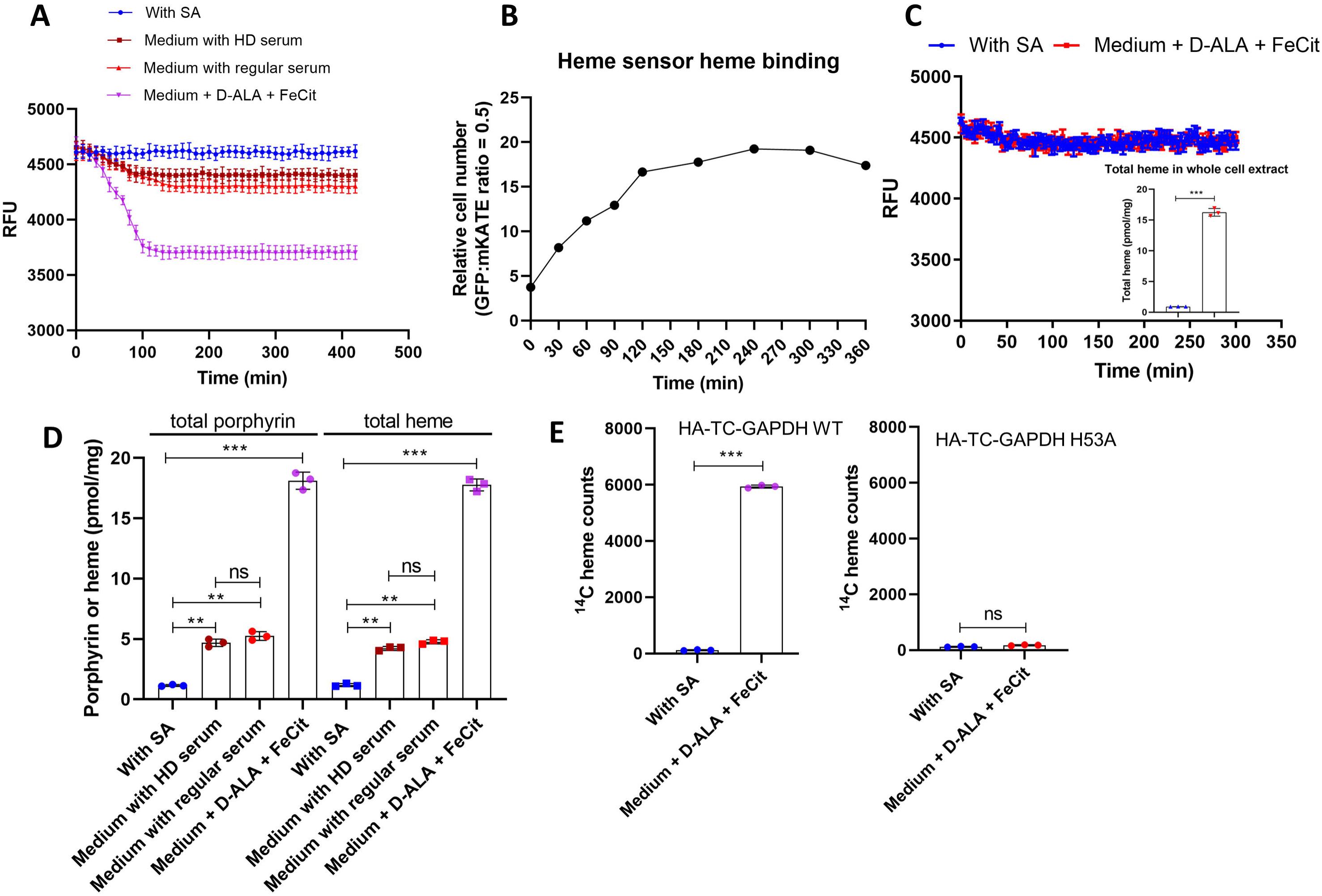
Heme binding by FlAsH-labeled HA-TC-hGAPDH expressed in living cells is indicated by quenching of the FlAsH fluorescence. HEK293T cells were cultured in a 96-well plate with SA to become heme deficient, were transfected to express HA-TC-hGAPDH or its H53A variant, FlAsH labeled, and then subject to the different culture conditions under continuous reading in a fluorescent plate reader. *Panel A-* Change in the cell FlAsH-HA-TC-hGAPDH fluorescence intensities with time after they were placed under the indicated culture conditions. *Panel B-* Change in the cell heme level with time after providing δ-ALA + Fe-cit to the cultures, as indicated by a fluorescent heme sensor protein expressed in the cells. *Panel C-* Fluorescence intensity versus time of cells expressing the FlAsH-labeled HA-TC-hGAPDH H53A variant, which has a defect in heme binding. Medium + SA or + δ-ALA and Fe-cit were given to the cells at time = 0. *Inset-* heme concentrations in the cell supernatants made at the end of the experiment. *Panel D-* Total porphyrin and heme concentrations in cell supernatants from Panel A made at the end of the experiment. *Panel E-* Radiolabel heme counts in IP pulldowns of supernatants from cells expressing HA-TC-hGAPDH WT or the H53A variant that were given ^14^C-δ-ALA + Fe-cit at time = 0 and then cultured for 300 min before lysis. Data represented as mean ± s.d. for n=3. **p<0.01 and ***p<0.001, student t-test in GraphPad Prism (v9).

### NO regulation of cellular heme allocation

We next utilized HA-TC-hGAPDH to investigate NO regulation of GAPDH-dependent intracellular heme allocation. We recently reported that very low levels of NO increased cell heme allocation to heme proteins, but above this range the positive effect of NO was quickly lost and could even reverse such that at higher levels there was a net loss of heme from some hemeproteins (17). We thus followed binding of mitochondrially-derived heme to HA-TC-hGAPDH in living HEK293T cells that were undergoing the low and high NO exposures. In most experiments, we co-expressed the heme protein IDO1, whose normal and NO-driven heme deliveries both depend on GAPDH and Hsp90 (7) and whose heme allocation displays the aforementioned hormetic response toward NO concentration (17). To generate precise NO exposure levels, we utilized the well-characterized slow-releasing NO donor molecule NOC-18 (t_1/2_ = 13 h under our conditions) as used in our previous study (17), and we investigated the impact of two lower NOC-18 concentrations (2.5 and 5 µM) known to promote cell heme allocation to IDO1 and one higher concentration (100 µM) known to cause heme loss from IDO1 (17).

We first studied the impact of these NOC-18 exposures on cells that expressed HA-TC-hGAPDH and IDO1 and were being cultured under normal conditions (i.e., they had normal resting levels of heme and could generate mitochondrial heme). Figure 3A shows FlAsH fluorescence traces collected after adding fresh media alone or media containing the different doses of NOC-18, along with a fluorescence trace collected for cells expressing HA-TC-hGAPDH and IDO1 under the fully heme-depleted condition to provide a comparative fluorescence signal intensity trace for heme-depleted HA-TC-hGAPDH. The relative fluorescence levels recorded at T = 0 indicate that HA-TC-hGAPDH contained more heme when it was expressed in the cells cultured normally versus in the heme-deprived cells, consistent with data in Fig. 2. This difference correlated to the levels of total porphyrin or total heme present in the cells under the two culture conditions (Fig. 3B). NOC-18 given at the two lower concentrations (2.5 and 5 µM) caused no discernable change in the proportion of heme-bound HA-TC-hGAPDH, as judged by the fluorescence signal intensity remaining unchanged over time during NOC-18 exposure (Fig. 3A), nor did these NOC-18 exposures change the final total porphyrin or heme levels present in the cells (Fig. 3B), despite their increasing cell heme allocation to IDO1 in a dose-dependent manner, as indicated by the gain in IDO1 enzymatic activity (Fig. 3C) and as consistent with our previous findings (17). Measures of the cell supernatant IDO1 activities after preincubation with either vehicle or added heme (6 µM) (17) showed that the 2.5 and 5 µM NOC-18 exposures caused the heme saturation level of IDO1 to increase from 15% to 40% and to 90% respectively, in the cells. By comparison, in the cells treated with 100 µM NOC-18, the level of heme in FlAsH-labeled HA-TC-hGAPDH increased *versus* time, as indicated by the gradual decrease in its fluorescence signal intensity versus time that was seen in this circumstance (Fig. 3A). Although there was no change to the final total porphyrin or heme levels in the cells cultured with 100 µM NOC-18 (Fig. 3B), there was a loss of IDO1 enzymatic activity and heme content under this circumstance (Fig. 3C and D), as we have observed previously (17). Western blot analyses showed that the HA-TC-hGAPDH or IDO1 showed similar expression levels among all the cell groups studied above and were not changed by the various experimental conditions (Fig. S5). Thus, the lower-level NO exposures stimulated cell heme allocation to IDO1 while keeping the level of heme-bound HA-TC-hGAPDH constant in the cells. In contrast, the higher NO exposure increased the proportion of heme-bound HA-TC-hGAPDH in the cells and caused a coincident loss of heme from IDO1. All these changes took place in the cells despite the NO exposures causing no change in their final total porphyrin or heme levels.

**Figure 3.**
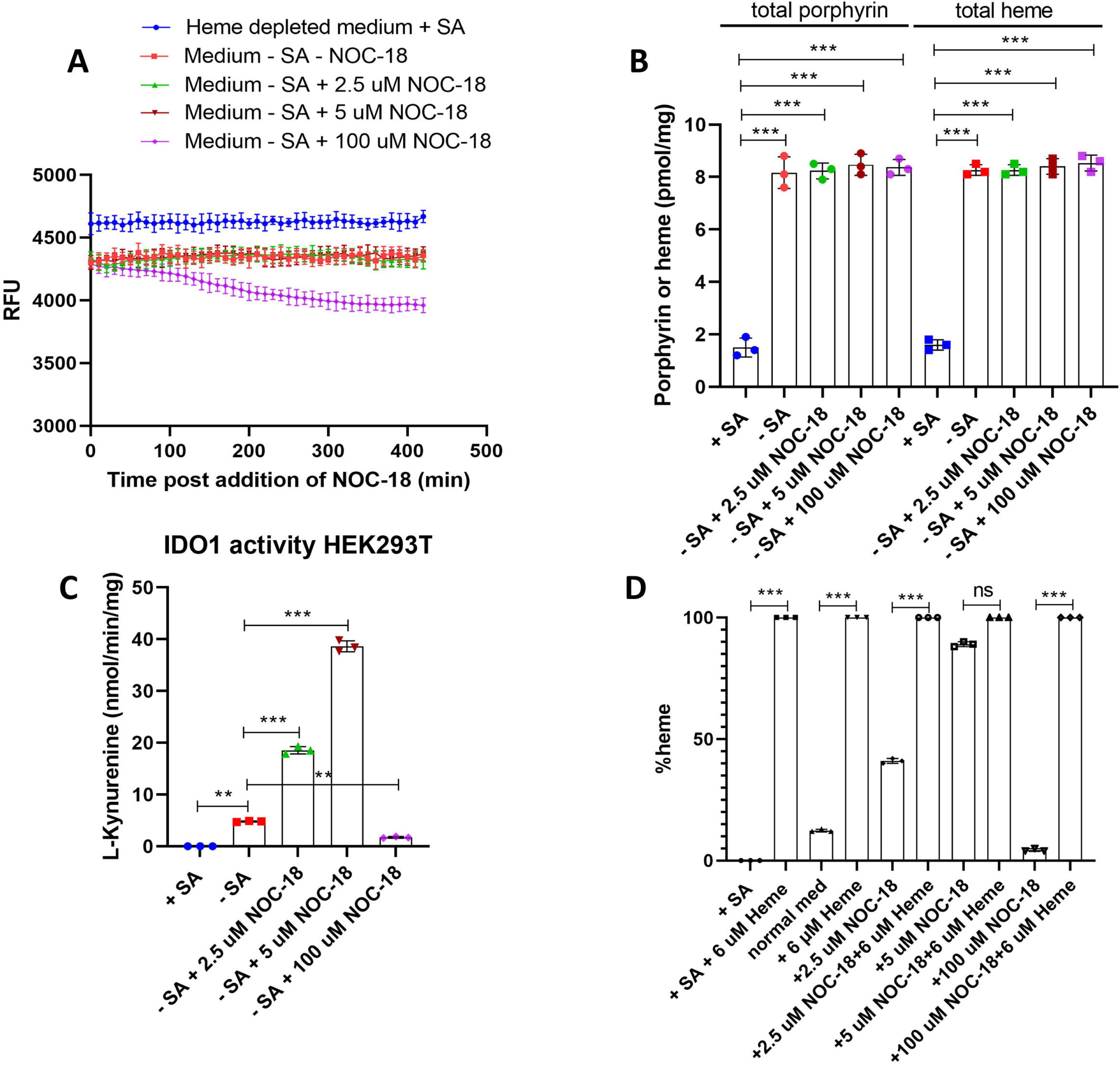
Effect of low or high NO exposure on heme levels in HA-TC-GAPDH and IDO1 in cells that maintained an active mitochondrial heme biosynthesis. HEK293T cells were cultured in a 96-well plate with or without SA as indicated, were transfected to express HA-TC-hGAPDH and IDO1, FlAsH labeled, and then subject to the indicated culture conditions under continuous reading in a fluorescent plate reader. *Panel A-* Change in the cell FlAsH-HA-TC-hGAPDH fluorescence intensities with time under the indicated conditions. *Panel B-* Total porphyrin and heme concentrations in cell supernatants from Panel A made at the end of the experiment. *Panel C-* IDO1 activity in supernatants of cells at the end of the experiment. *Panel D-* Heme saturation level of IDO1 that was achieved under each culture condition, determined by comparing the activities of each supernatant after they had or had not undergone incubation with added heme. Data represented as mean ± s.d. for n=3. **p<0.01 and ***p<0.001, student t-test in GraphPad Prism (v9).

Because the study above utilized cells whose mitochondrial heme synthesis remained active during the NOC-18 exposures, we wondered how having access to new mitochondrial heme synthesis may have impacted the results. We therefore repeated the study using cells whose mitochondrial heme production was halted by adding SA just at the point of the NOC-18 additions. SA is reported to inhibit ALAS activity and heme synthesis in cells within minutes (27,28). In this circumstance, the cells would be left with the amount of heme that they presently contain, which then becomes subject to possible redistribution by the NOC-18 treatments. Results are shown in Fig. 4. In the absence of added NOC-18, the cells maintained a steady level of heme-bound HA-TC-hGAPDH despite their mitochondrial heme synthesis having been turned off by the SA. The addition of NOC-18 at 2.5 and 5 µM now caused a time-dependent loss of heme from HA-TC-hGAPDH in the cells, as indicated by the rising fluorescence traces versus time (Fig. 4A). In the case of the 5 µM NOC-18 treatment, this led to a near complete loss of heme from HA-TC-hGAPDH by the end of the time course, as judged by the fluorescent trace intensity nearing the high level trace that was recorded for heme-depleted HA-TC-hGAPDH in the heme-depleted cells in the same plate (Fig. 4A). In contrast, the fluorescent traces indicate that the 100 µM NOC-18 exposure caused cells to gradually load existing heme onto their HA-TC-hGAPDH, along with causing a coincident loss of heme from IDO1 (Fig. 4C and D). The final total porphyrin or heme levels in the cells were not altered by the NOC-18 treatments (Fig. 4B). The NOC-18 treatments caused dose-related changes in the IDO1 activity as was seen before in cells with active mitochondrial heme synthesis (Fig. 4C). However, in the current circumstance there was less heme incorporation into IDO1 in response to the 2.5 and 5 µM NOC-18 exposures, such that it reached only a 25 and 55% heme saturation level, respectively (Fig. 4D), compared to IDO1 reaching a 90% heme saturation level in response to 5 µM NOC-18 in cells whose mitochondrial heme synthesis remained active (compare Fig. 3D). Western blot analyses showed that the protein expression levels of HA-TC-hGAPDH or IDO1 proteins were similar among all the cell groups and were not changed by the various experimental conditions (Fig. S6). Thus, the NOC-18 treatments caused cells with a halted mitochondrial heme production to redistribute their existing heme into or out of IDO1 in a similar manner, but with two notable differences: (a) the redistribution of existing heme into IDO1 now correlated with a proportional and nearly complete loss of heme from HA-TC-hGAPDH, and (b) the amount of existing heme available for redistribution appeared to be insufficient to saturate the IDO1 enzyme, despite the cells containing total final heme levels that were similar to those in cells whose mitochondrial heme synthesis was not blocked (compare Panel B in Figs. 3 and 4).

**Figure 4.**
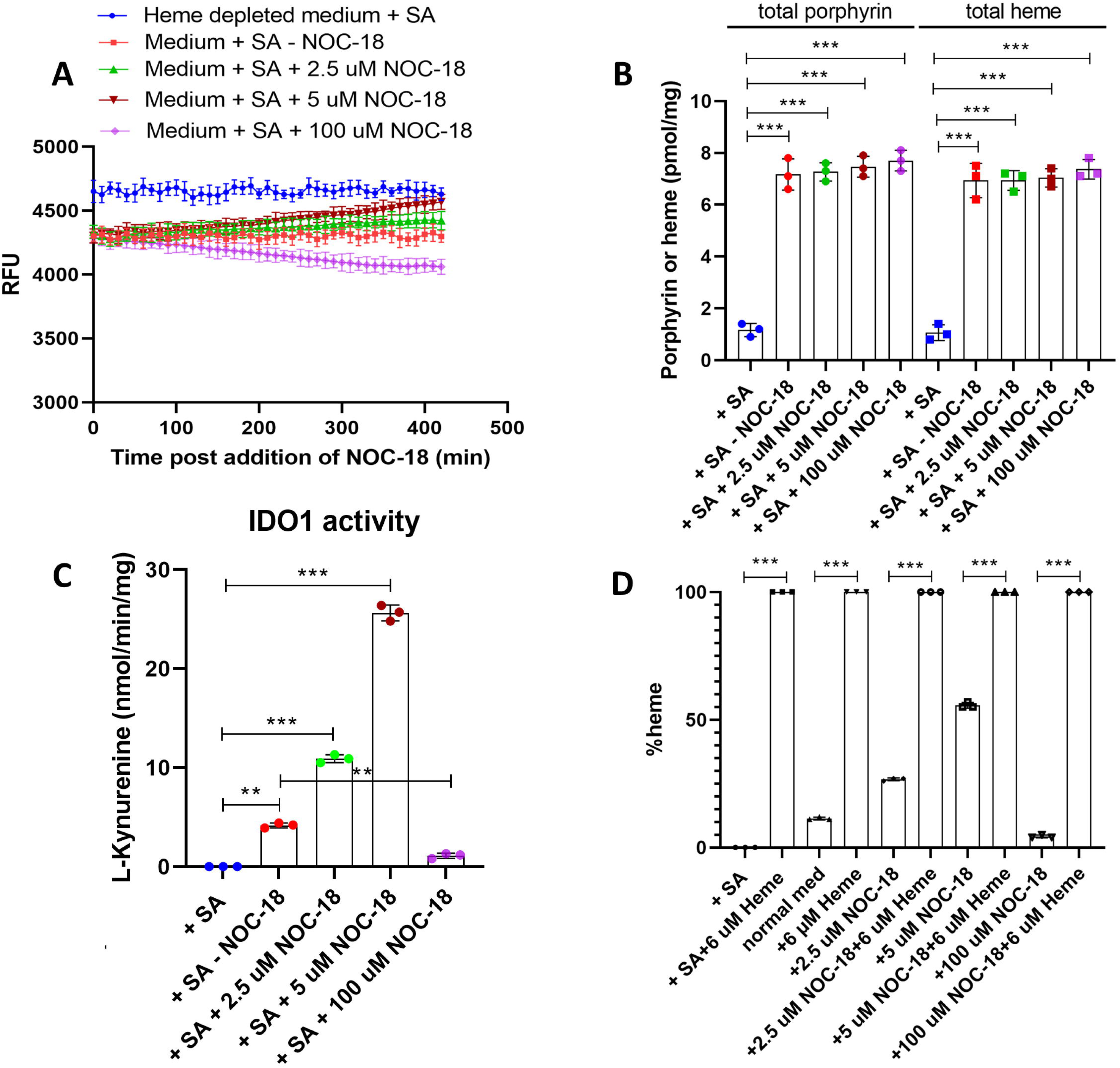
Effect of low or high NO exposure on heme levels in HA-TC-hGAPDH and IDO1 in cells whose mitochondrial heme biosynthesis was blocked at the time of NOC-18 addition. HEK293T cells were cultured in a 96-well plate, transfected to express HA-TC-hGAPDH and IDO1, FlAsH labeled, and then given NOC-18 along with SA to stop mitochondrial heme synthesis just before readings commenced. *Panel A-* Change in the cell FlAsH-HA-TC-hGAPDH fluorescence intensities with time under the indicated conditions. *Panel B-* Total porphyrin and heme concentrations in cell supernatants from Panel A measured at the end of the experiment. *Panel C-* IDO1 activity in supernatants from cells cultured under each condition. *Panel D-* Heme saturation level of IDO1 that was achieved under each culture condition, determined as in Fig 3D. Data represented as mean ± s.d. for n=3. **p<0.01 and ***p<0.001, student t-test in GraphPad Prism (v9).

To explore this further, we repeated the experiments using HEK293T cells that had now been pre-incubated 48 h with δ-ALA and Fe-cit to increase their total heme level by 3 to 4 times (see Fig. 2) before doing the NOC-18 exposures. The cells were transfected to express HA-TC-hGAPDH and IDO1 as before and had new mitochondrial heme synthesis blocked by adding SA at the point of NOC-18 additions. Figure 5A shows that there was a much greater difference in the initial fluorescence intensity values at T = 0 between cells precultured with δ-ALA and Fe-cit and for the heme-depleted cells, indicating the pre-incubated cells contained a higher proportion of heme-bound HA-TC-hGAPDH, as expected, and in line with results from Fig 2 and with the cells having 3 times higher total porphyrin and heme levels (compare Figs. 4B and 5B). The addition of NOC-18 at 2.5 and 5 µM caused a gradual and dose-dependent loss of existing heme from HA-TC-hGAPDH (Fig. 5A) as indicated by the gain in the fluorescence signal intensities, which equilibrated to new equilibrium levels after about 350 min of low NOC-18 exposures, but did not reach the higher level seen for HA-TC-hGAPDH that was expressed in the heme-depleted control cells (Fig. 5A). This indicated that some existing heme remained bound in HA-TC-hGAPDH after the two low NOC-18 exposures. In comparison, the 100 µM NOC-18 treatment again caused cells to gradually reallocate existing heme onto HA-TC-hGAPDH until a new equilibrium was reached by around 250 min (Fig. 5A). None of the treatments caused changes in the total cell porphyrin or heme levels (Fig. 5B) nor impacted the HA-TC-hGAPDH or IDO1 protein expression levels, which were similar among the groups (Fig. S7). Regarding IDO1, its activity increased about 3-fold just due to the δ-ALA plus Fe-cit preincubation alone (compare data in Figs. 4C and 5C) and increased a further 2.5 and 5-fold in response to the 2.5 and 5 µM NOC-18 exposures, respectively (Fig. 5C). These IDO1 activity changes were tied to a rise in its heme level from 10 to 20% saturated due to the δ-ALA plus Fe-cit treatment alone (compare data in Figs. 4D and 5D) and to further increases in its heme content in response to the subsequent 2.5 and 5 µM NOC-18 exposures from 20% heme saturated to 50 and 100%, respectively (Fig. 5D). In comparison, the 100 µM NOC-18 treatment again caused IDO1 to lose both activity and its existing heme (Fig. 5C and D) despite the cells having a 3 times greater than normal total heme content under this circumstance (Fig. 5B).

**Figure 5.**
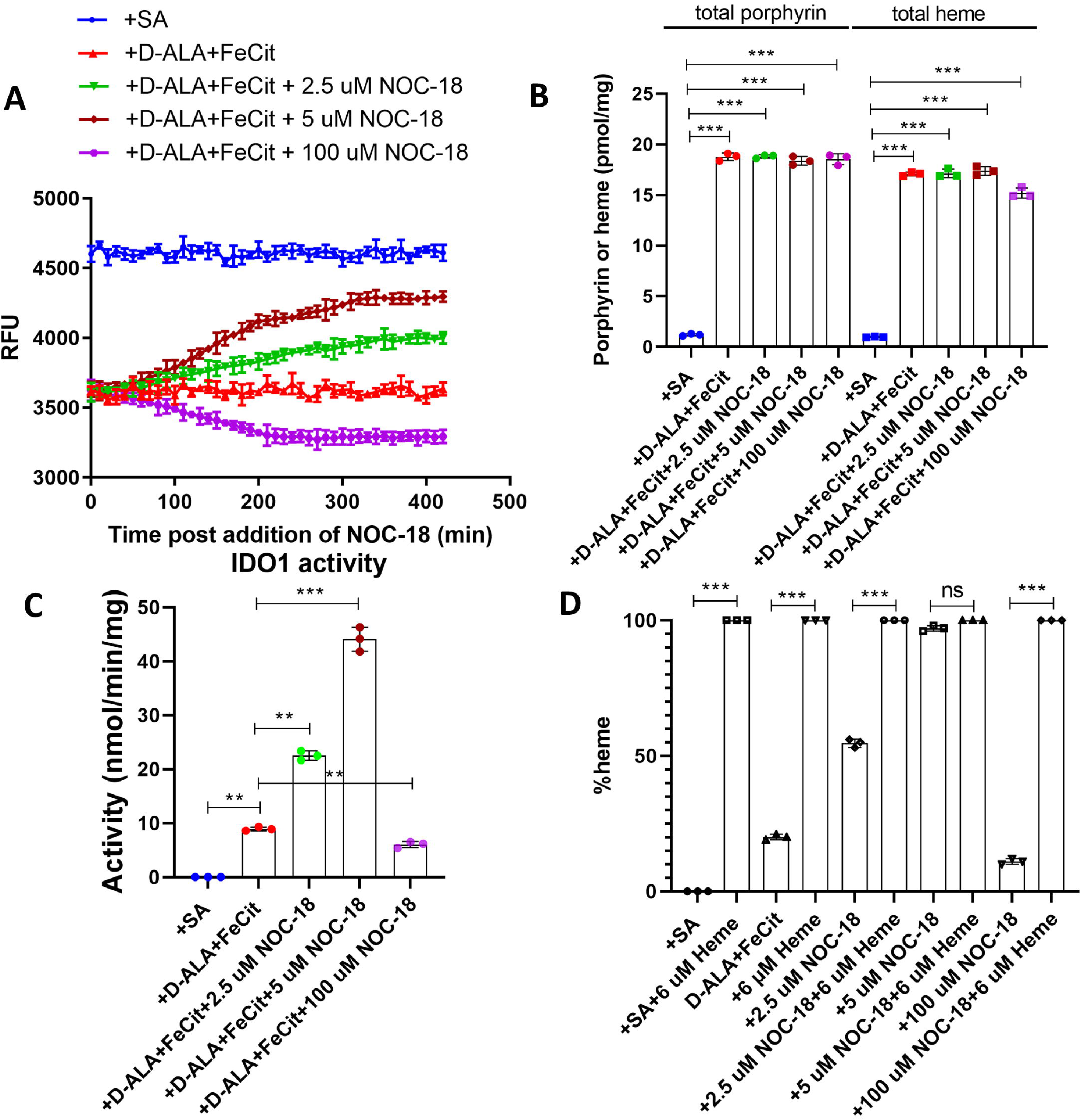
Effect of low or high NO exposure on heme levels in HA-TC-hGAPDH and IDO1 in cells whose mitochondrial heme biosynthesis had been stimulated prior to being blocked at the time of NOC-18 addition. HEK293T cells were pre-cultured with δ-ALA + Fe-cit to increase their heme level, transfected to express HA-TC-hGAPDH and IDO1, FlAsH labeled, and NOC-18 was given along with SA to stop mitochondrial heme synthesis just before readings commenced. *Panel A-* Change in the cell FlAsH-HA-TC-hGAPDH fluorescence intensities with time under the indicated conditions. *Panel B-* Total porphyrin and heme concentrations in cell supernatants from Panel A at the end of the experiment. *Panel C-* IDO1 activity in supernatants of cells cultured under each condition. *Panel D-* Heme saturation level of IDO1 that was achieved under each culture condition, determined as in Fig 3D. Data represented as mean ± s.d. for n=3. **p<0.01 and ***p<0.001, student t-test in GraphPad Prism (v9).

### NO-driven reallocation of GAPDH-heme requires that IDO1 acceptor be present and can accept heme

The reciprocal change in the levels of existing heme in HA-TC-hGAPDH and IDO1 that we observed upon low NO exposure suggested the possibility of heme exchange occurring between the two proteins. To investigate, we repeated the experiment with δ-ALA and Fe-cit preincubated cells that were transfected only to express HA-TC-hGAPDH. As before, the cells had their mitochondrial heme synthesis blocked by adding SA at the point of NOC-18 addition. In this circumstance (Fig. 6A) the 2.5 and 5 µM NOC-18 exposures caused no discernable loss of existing heme from HA-TC-hGAPDH, quite unlike the significant loss that occurred when IDO1 was co-expressed in the cells under the same circumstances (compare to Fig. 5A). The 100 µM NOC-18 treatment still caused existing heme to load onto HA-TC-hGAPDH (Fig. 6A) but did so to a lesser extent than what was observed when IDO1 was co-expressed in the cells (compare to Fig. 5A). The total porphyrin or heme levels remained relatively constant among the cell groups (Fig. 6B) and the HA-TC-hGAPDH expression levels were similar (Fig. S8). Thus, the extent to which low NO drove the loss of existing heme from HA-TC-hGAPDH depended heavily on IDO1 being co-expressed in the cells, consistent with low NO driving a specific exchange of heme from HA-TC-hGAPDH to IDO1.

**Figure 6.**
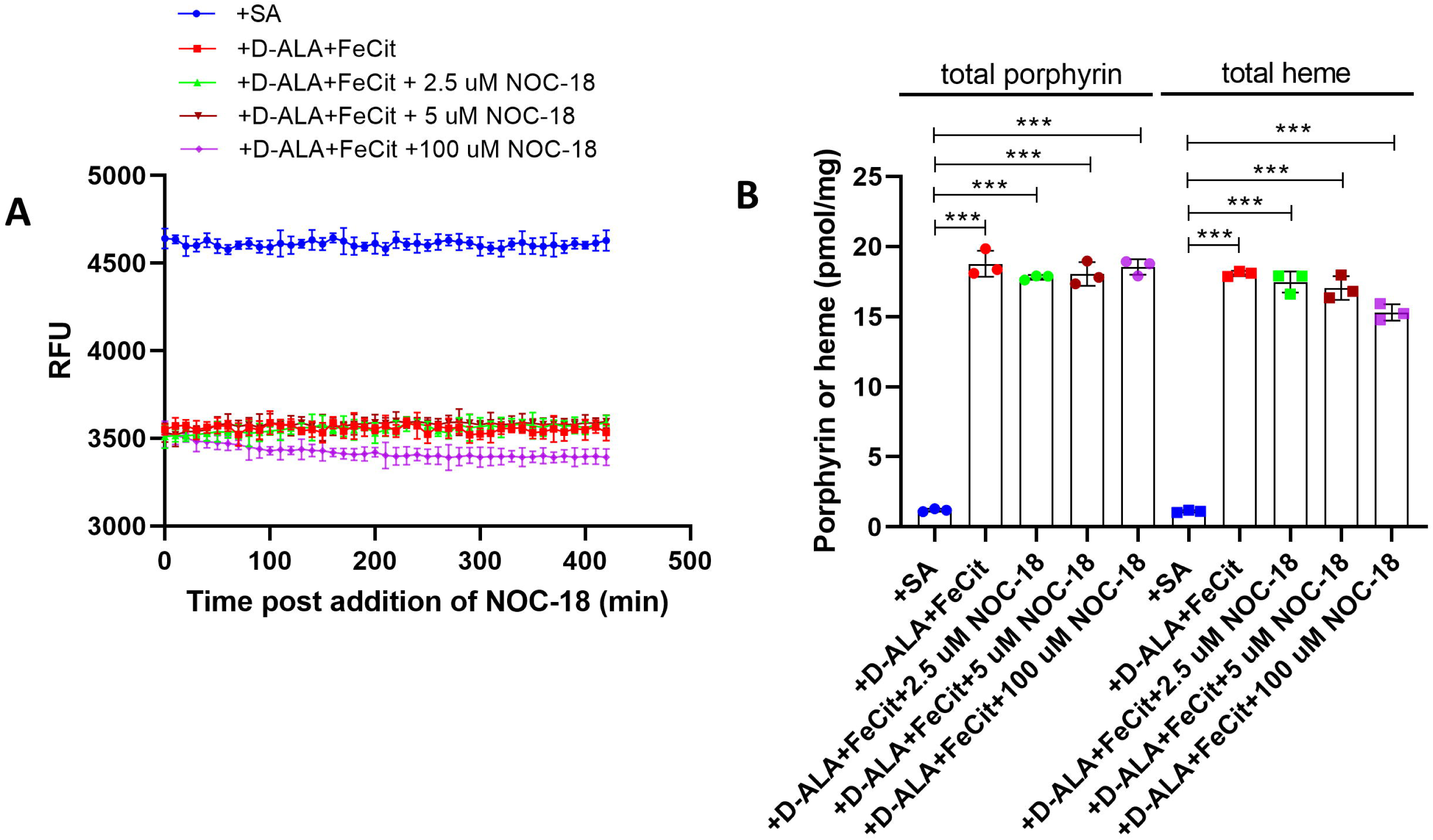
NO-driven reallocation of heme from HA-TC-hGAPDH requires IDO1 co-expression. HEK293T cells were precultured with δ-ALA + Fe-cit to increase their heme level, were transfected to express HA-TC-hGAPDH alone, FlAsH labeled, and NOC-18 was given along with SA to stop mitochondrial heme synthesis just before readings commenced. *Panel A-* Change in the cell FlAsH-HA-TC-hGAPDH fluorescence intensities with time under the indicated conditions. *Panel B-* Total porphyrin and heme concentrations in cell supernatants from Panel A at the end of the experiment. Data represented as mean ± s.d. for n=3. ***p<0.001, student t-test in GraphPad Prism (v9).

To probe further we examined how the NO-driven changes in the existing heme levels in HA-TC-hGAPDH and in IDO1 might be influenced by the cell chaperone Hsp90. Our previous studies showed that Hsp90 is bound to the heme-free apo-IDO1 in cells and the ATP-ase function of Hsp90 enables heme insertion into apo-IDO1 both during its normal maturation (7) and during NO-driven heme allocation to IDO1 (17). Our HA-TC-hGAPDH construct also provided an opportunity to determine if Hsp90 might be involved in the upstream delivery of mitochondrial heme to GAPDH. To test this possibility, HEK293T cells were transfected to express HA-TC-hGAPDH and then given buffer with or without the Hsp90 inhibitor radicicol for 6 h before stimulating cell mitochondrial heme production by adding δ-ALA plus Fe-cit. Figure 7A shows that the Hsp90 inhibitor had no impact on the ability of δ-ALA plus Fe-cit to increase mitochondrial heme allocation to HA-TC-hGAPDH, and this was confirmed in replica experiments that utilized ^14^C-δ-ALA and Ab pulldown of GAPDH (Fig. 7B). Thus, we conclude that Hsp90 is not involved in the upstream allocation of mitochondrial heme to GAPDH.

**Figure 7.**
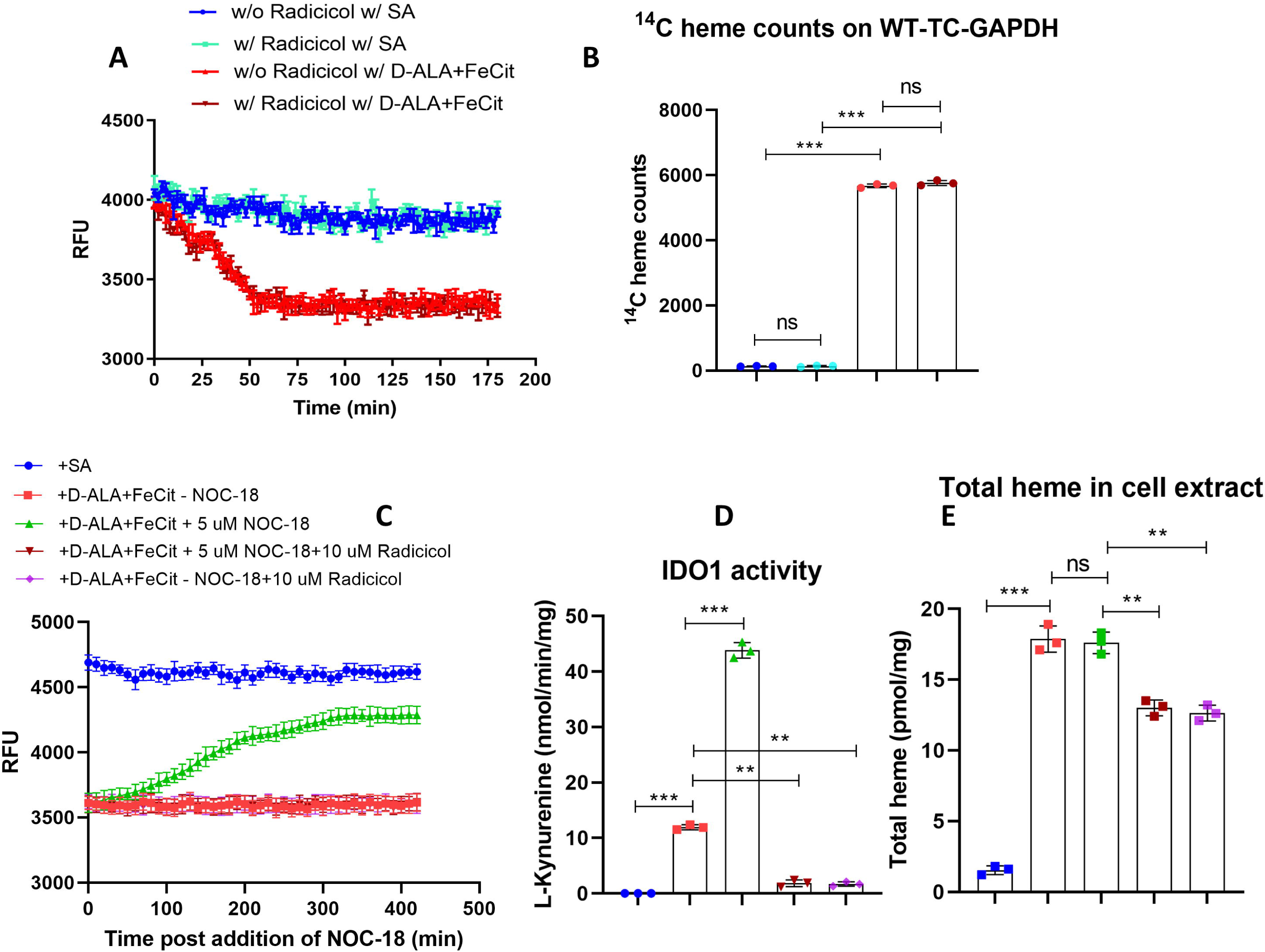
NO-driven reallocation of heme from HA-TC-GAPDH requires that IDO1 can accept the heme. HEK293T cells that did or did not undergo δ-ALA + Fe-cit pre-culture were transfected to express HA-TC-hGAPDH and IDO1, FlAsH labelled, Hsp90 inhibitor radicicol was added in some cases, and then δ-ALA + Fe-cit and/or 5 µM NOC-18 were given along with SA to stop mitochondrial heme synthesis just before readings commenced. *Panel A-* Impact of radicicol on HA-TC-hGAPDH mitochondrial heme acquisition in response to δ-ALA + Fe-cit addition. *Panel B-* Replica experiment showing the effect of radicicol on ^14^C heme counts accumulated in HA-TC-hGAPDH pulldowns from cells cultured for 180 min with ^14^C-δ-ALA and Fe-cit. *Panel C-* Effect of radicicol on the ability of NO to drive heme off HA-TC-hGAPDH in cells preincubated with δ-ALA + Fe-cit. *Panel D-* IDO1 activity indicated by L-Kynurenine production by cell supernatants from Panel C. *Panel E-* Total heme levels in cell supernatants from panel C. Data represented as mean ± s.d. for n=3. **p<0.01 and ***p<0.001, student t-test in GraphPad Prism (v9).

Armed with this knowledge, we next investigated if Hsp90 inhibition might impact the NO-driven reallocation of existing heme from HA-TC-hGAPDH to IDO1. HEK293T cells that were pre-incubated with δ-ALA plus Fe-cit and were transfected to express HA-TC-hGAPDH and IDO1 had their mitochondrial heme synthesis blocked by adding SA at the point of 5 µM NOC-18 addition. Figure 7C shows that radicicol completely prevented the loss of existing heme from HA-TC-hGAPDH that normally took place in response to 5 µM NOC-18 under this circumstance. Moreover, this correlated with heme allocation to IDO1 being blocked, as was indicated by IDO1 activity measures (Fig. 7D). Cell total heme levels remained similar among the groups although they were 30% lower in the radicicol-treated cells (Fig. 7E), and the expression level of HA-TC-hGAPDH was similar throughout (Fig. S9). Thus, NO-driven reallocation of the existing heme in HA-TC-hGAPDH depended on an acceptor heme protein like IDO1 being both present and capable of accepting the heme from HA-TC-hGAPDH.

GAPDH heme allocation to IDO1 is reversible depending on the NO concentration Finally, to explore the possibility that NO can induce a concentration-dependent, reversible exchange of existing heme between HA-TC-hGAPDH and IDO1; we studied the impact of consecutive cell exposure to 5 and 100 µM NOC-18. We utilized HEK293T cells that were preincubated with δ-ALA plus Fe-cit and were expressing HA-TC-hGAPDH and IDO1, and blocked mitochondrial heme synthesis by giving SA at the point of the initial 5 µM NOC-18 addition. Figure 8A and B show that the 5 µM NOC-18 exposure caused the cells to reallocate existing heme from HA-TC-hGAPDH into IDO1 over the first 300 min, as expected and observed previously. Exposing the cells at this point to 100 µM NOC-18 caused a time-dependent reversal in the heme reallocation, such that the existing heme now loaded out of IDO1 and back into HA-TC-hGAPDH over the next 300 min (Fig. 8A and B). In the absence of NOC-18 the levels of existing heme bound in HA-TC-hGAPDH or IDO1 stayed constant throughout the entire 600 min period (Fig. 8A and B), and the NOC-18 exposures did not impact the total cell heme levels (Fig. 8C) or the expression levels of either protein (Fig. S10). Thus, NO-driven reallocation of existing heme between HA-TC-hGAPDH and IDO1 was reciprocal and reversible, going in either direction depending on the level of NO exposure.

**Figure 8.**
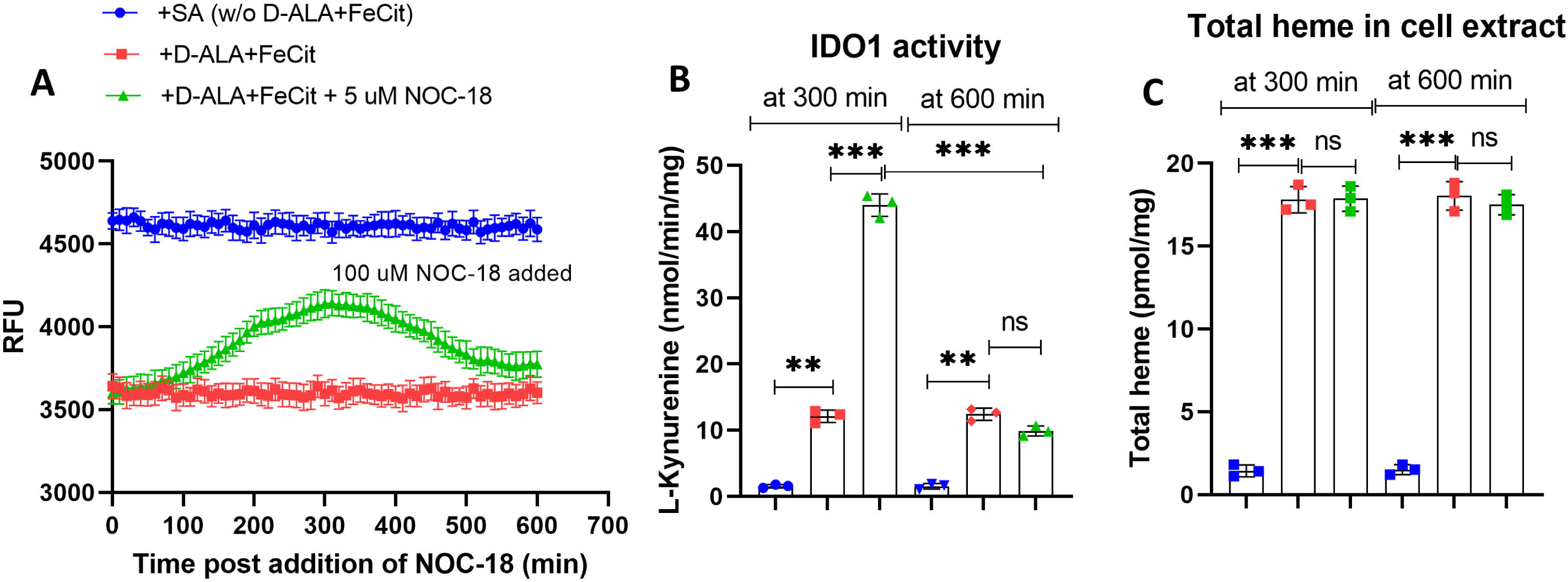
Existing heme in HA-TC-hGAPDH and IDO1 reallocates in opposite directions depending on the NO exposure level. HEK293T cells were pre-cultured with SA or δ-ALA + Fe-cit as indicated, transfected to express HA-TC-hGAPDH and IDO1, FlAsH labeled, and given buffer or 5 µM NOC-18 along with SA to stop mitochondrial heme synthesis just before readings commenced. At 300 min NOC-18 was increased to 100 µM and the readings continued. *Panel A-* Change in FlAsH-HA-TC-hGAPDH fluorescence in cells cultured under the indicated conditions. *Panel B-* IDO1 activity as indicated by L-Kynurenine production by supernatants of cells cultured under the indicated conditions and harvested at 300 or 600 min. *Panel C-* Total heme levels in supernatants of the cells from panel B. Data represented as mean ± s.d. for n=3. **p<0.01 and ***p<0.001, student t-test in GraphPad Prism (v9).

## Discussion

GAPDH has various functions in biology including its chaperoning mitochondrially-generated heme for allocation in mammalian cells (13). The TC-hGAPDH reporter construct described here behaved like native GAPDH regarding its physical, heme binding, and enzymatic properties, and thus upon its FlAsH labeling could serve as a probe to report on GAPDH heme binding in living mammalian cells in real time using a conventional 96-well fluorescent plate reader. This new approach to studying intracellular heme trafficking sheds light on several aspects, as discussed below.

### GAPDH heme binding in cells

The fluorescence quenching behavior of purified FlAsH-labeled TC-hGAPDH upon heme binding indicated its heme affinity was similar to wild-type hGAPDH and that it contained two high-affinity heme binding sites per tetramer that displayed no apparent binding co-operativity. When two heme bound to the FlAsH-TC-hGAPDH in its purified form or when it was expressed in HEK293T cell supernatants it reduced its fluorescence emission intensity by approximately 44 ± 7% (n = 3) and 65 ± 2% (n = 5), respectively. If we assume the latter percentage decrease coincides with full heme loading onto TC-hGAPDH (i.e., 2 heme per tetramer) when it is expressed in mammalian cells, it can be used to estimate the HA-TC-hGAPDH heme saturation level in cells, based on the differences in the fluorescence emission intensities that we observed for FlAsH-labeled HA-TC-hGAPDH when it was expressed in fully heme-deficient (SA-treated) cells versus in cells that were cultured under other conditions. For example, a comparison of the fluorescence traces in Fig. 2 suggests that in HEK293T cells cultured with normal media and serum their HA-TC-hGAPDH contained approximately 0.20 heme per tetramer, which, assuming HA-TC-hGAPDH equilibrates with the cellular pool of GAPDH regarding heme binding, suggests that most of the GAPDH tetramers in the cells were devoid of heme under this culture condition. In comparison, in the cells that had been cultured with the mitochondrial heme precursors δ-ALA and Fe-cit to increase their total heme level by 3-fold, the fluorescent traces in Fig. 2 indicate that after equilibrium the HA-TC-hGAPDH contained 0.65 heme per tetramer, suggesting that a little more than half of the cell GAPDH would contain 1 heme per tetramer in this circumstance. If we further assume a GAPDH tetramer concentration of 100 µM exists in mammalian cells (29), and there is no GAPDH compartmentalization with regard to heme chaperoning, this suggests that in the cells cultured under normal conditions or that were incubated with the heme precursors δ-ALA and Fe-cit, the GAPDH-heme complex was present at intracellular concentrations of 20 and 65 µM respectively. Despite all the assumptions, these estimates provide a first glimpse of the possible range of heme binding levels that HA-TC-hGAPDH (and by extension, GAPDH) may adopt in mammalian cells, and imply that when HEK293T cells are grown under normal culture conditions the proportion of their GAPDH tetramers that contain heme is relatively low. Such sub-stoichiometric heme binding is what one might expect for a protein that is expressed constitutively at a relatively high concentration (29,30) and functions to chaperone intracellular heme, and so must maintain an intermediate affinity. It is also consistent with the concept that mammalian cells maintain a natural heme deficit that prevents their heme proteins from becoming fully heme-saturated (15), as was demonstrated here for IDO1 (only 15% heme saturated under normal culture conditions) and previously for other heme proteins as well (15). In future it will be interesting to see if the percentage of heme-bound GAPDH differs in various cell types or animal species and to determine how its level is regulated. One can also consider how such levels of GAPDH heme binding might influence its other cellular functions (31). Regardless, our current findings provide a first indication of the heme binding range in which GAPDH operates to enable intracellular heme trafficking in HEK293T cells.

We also saw that the level of heme-bound HA-TC-hGAPDH remained steady in cells over 6.5 h experimental observation periods, irrespective of whether we allowed cell mitochondrial heme biosynthesis to remain active or to have been inhibited at the start of the time course, or despite our inhibiting new protein synthesis during the observation time with cycloheximide. This held under all conditions of culture that we examined (cells that were heme-deficient, or had normal or supranormal heme levels), and correlated with the cells maintaining constant levels of total heme. Thus, cells conserved both their existing total heme and their HA-TC-hGAPDH-bound heme and did not need active mitochondrial heme biosynthesis to do so within the 6.5 h time period. This is consistent with two to three days of culture being required for cells to become heme-depleted after inhibiting their mitochondrial heme synthesis with SA (24), and it implies that under normal culture conditions loss of existing GAPDH by protein turnover or of GAPDH heme via its catabolism by heme oxygenases or efflux through cell membrane heme exporters is negligible within a 6.5 h time period.

### Regulation of cell heme allocation by low NO

NO displays a remarkable hormetic effect on cell heme allocation to IDO1 and TDO, where low NO exposure boosts the heme allocations while higher NO levels no longer support allocation and cause heme loss (15). In our current experiments we chose to co-express IDO1 as the heme protein target for HA-TC-hGAPDH, because it obtains mitochondrial heme via GAPDH in an Hsp90-dependent process (7) and its NO-driven heme allocation also depends on GAPDH and Hsp90 acting in the same ways (17).

When cells cultured under normal conditions maintained an active mitochondrial heme biosynthesis, the low-level NO exposures stimulated heme allocation to IDO1 despite the level of heme-bound HA-TC-hGAPDH remaining constant. In this circumstance, the IDO1 heme content increased from 10 to 90%. In comparison, when cell mitochondrial heme synthesis was inhibited during the low NO exposures, reallocation of the existing cell heme to IDO1 still occurred but was now accompanied by a near total loss of heme from HA-TC-hGAPDH, and in this circumstance the increase in IDO1 heme content was blunted and rose to only 55% heme-saturated. These findings imply: (a) The existing level of GAPDH-bound heme that is present in cells cultured under normal conditions can become limiting when low NO exposure promotes heme reallocation from GAPDH to a client protein like IDO1, if the client is expressed in sufficient quantity. (b) When cell mitochondrial heme synthesis is active, the NO-driven reallocation of existing heme from HA-TC-hGAPDH to the IDO1 is accompanied by a coincident resupply of new mitochondrial heme to HA-TC-hGAPDH such that its level of bound heme remains constant, and this resupplying of new mitochondrial heme to GAPDH enables a fuller heme allocation to IDO1. Thus, a feedback regulation likely exists that can sense GAPDH heme loss and rebalance it by increasing the synthesis and/or provision of mitochondrial heme to GAPDH. This feedback loop may be related to the coupling that has been observed for example between the level of mitochondrial heme production and the amount of hemoglobin protein being actively expressed in erythrocytes (32,33), and further investigation is warranted.

As noted, in the mitochondrial-inhibited cells low NO exposures still drove significant reallocation of existing cell heme to IDO1 despite it resulting in a near-complete loss of heme from HA-TC-hGAPDH (and by extension, from GAPDH). Mechanistically, this implies that low NO acts on the heme reallocation pathway at or downstream from the existing GAPDH-heme species in cells, rather than it acting by enhancing “upstream” heme loading onto GAPDH. However, we saw that low NO did not promote heme reallocation from GAPDH simply by lowering its binding affinity so that its heme dissociated in a random manner into the cytosol. This was demonstrated by low NO only causing significant reallocation of heme from HA-TC-hGAPDH when an acceptor heme protein like IDO1 was co-expressed in the cells, and moreover, only when the co-expressed IDO1 could incorporate the heme. Thus, cells displayed a remarkable control over their GAPDH heme provision in response to low NO. Overall, the findings suggest a heme transfer mechanism that likely relies on direct GAPDH-client protein contact instead of a more random process where heme in GAPDH dissociates and travels through the cell to final targets, primarily based on thermodynamic considerations. It will now be interesting to test the importance and details of such GAPDH-client protein interactions in the heme transfer process.

### High NO exposure and a reversal of GAPDH heme allocation

A higher level NO exposure (100 µM NOC-18) impacted heme allocation in a remarkably different way. It caused an increase in heme-bound HA-TC-hGAPDH, rather than the decrease that was seen with the low NO exposures. This increase occurred irrespective of whether cell mitochondrial heme synthesis was active or not, and it was independent of IDO1 co-expression, although without IDO1 the increase in heme-bound HA-TC-hGAPDH in the cells was significantly muted. Indeed, higher NO exposure appeared to cause the heme in IDO1 to reallocate to HA-TC-hGAPDH. This was most clearly demonstrated in the experiment where cells were sequentially exposed to a low then higher concentration of NOC-18. The low NO exposure caused cells to reallocate their existing HA-TC-hGAPDH heme to IDO1, and then the subsequent higher NO exposure caused the same cells to reallocate the existing heme in IDO1 back to HA-TC-hGAPDH. To our knowledge, this is the first evidence that heme allocation between two proteins can be reversible in living cells, as directed by NO in a concentration-dependent manner. This also reveals a way that cells may conserve their existing heme even after it has been allocated to a “final stage” heme protein like IDO1, namely through an NO-driven mechanism of heme banking on GAPDH. Whether this behavior is general or is specific to IDO1 remains to be examined, but precedent supports NO may have a general ability to drive heme exchange between proteins, at least in vitro (34,35). In any case, our findings highlight the ability of NO to drive heme reallocation between GAPDH and an acceptor protein in a forward or reverse direction depending on its concentration, thus revealing a new way that NO can impact the activities and functions of heme proteins inside the cell.

### Conclusions

Our findings are summarized in Fig. 9. A picture is emerging where under normal culture conditions, mitochondrial heme is allocated such that GAPDH and heme proteins are only partly heme-saturated in cells (15). Within this context, a balance is maintained between mitochondrial heme production and export, the heme level in GAPDH, and the heme level in a GAPDH-dependent client protein like IDO1. A low NO exposure on its own was a potent stimulus of cell heme allocation to IDO1, and the cells increased their flux of mitochondrial heme to GAPDH in kind such that its heme level remained constant in cells undergoing the low NO exposure. In cells whose mitochondrial heme synthesis had been inhibited at the point of low NO exposure, the NO still promoted reallocation of the existing heme in GAPDH to IDO1, but this caused a near total loss of the heme bound in GAPDH. In contrast, higher NO exposure drove the existing heme in IDO1 back onto GAPDH. Overall, this suggests a dynamic equilibrium exists between mitochondria, GAPDH, and IDO1 heme allocation, with NO being capable of driving the equilibrium between GAPDH and IDO1 in either direction according to its concentration. In the future, it will be important to examine the universality of these findings, particularly in cells that naturally express other heme proteins whose heme deliveries are GAPDH-dependent and can change in response to NO (17,19,21). Because these impacts of NO on GAPDH-dependent cell heme allocation occur within its physiologic range (19,36,37), these processes stand to influence heme protein functions in physiology and disease and should be further investigated.

**Figure 9.**
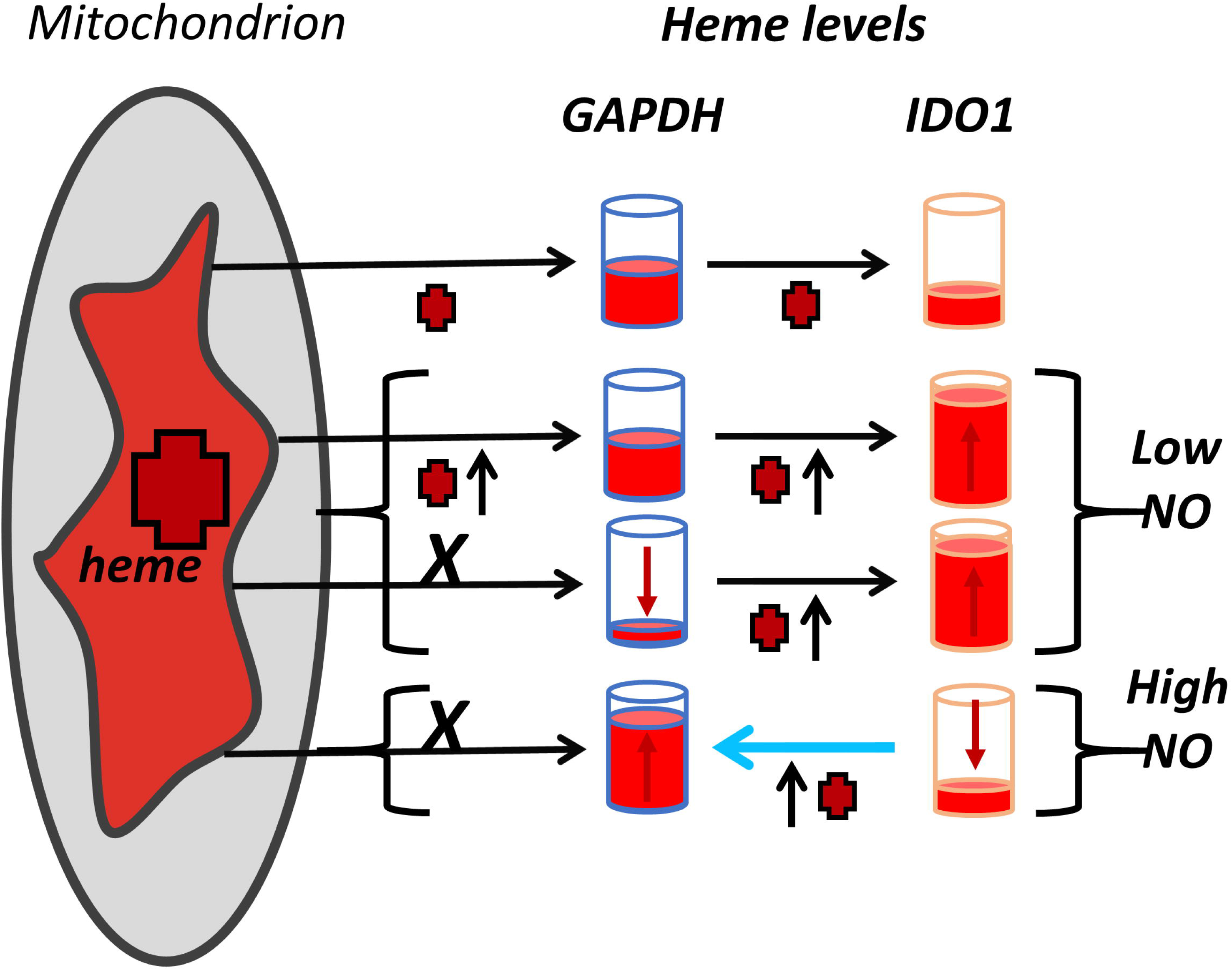
Cell allocation of mitochondrial heme to GAPDH and its client heme protein IDO1 and the impacts of low and high NO exposure. Heme is indicated by red crosses. Heme levels in GAPDH and IDO1 are indicated by the height of red liquid in the cylinders, which are quantitative for IDO1 but only relative for GAPDH for ease of illustration. Horizontal arrows indicate direction of heme flow, short vertical arrows indicate the change in heme flow or content. *Uppermost horizontal arrow set-* Under normal culture conditions cell mitochondrial heme distributes such that GAPDH and IDO1 are partially saturated with heme. *Middle two horizontal arrow sets-* A low-level NO exposure causes heme to shift from GAPDH to IDO1. When cell mitochondria can generate heme, they increase its provision to GAPDH so that the level of GAPDH-heme remains constant. If mitochondrial heme synthesis is blocked (*X*), the NO-driven heme transfer to IDO1 still occurs but depletes GAPDH of its heme. *Lowest horizontal arrow set-* Higher NO exposure causes the heme in IDO1 to transfer back to GAPDH (blue arrow).

## Experimental procedures

### Materials

[^14^C]-δ-ALA (1 µCi/µl) was purchased from ChemDepo Inc. All other reagents used were purchased from Sigma unless otherwise mentioned.

### Insertion of the tetra cysteine (TC) tag into human GAPDH gene

The structure of human GAPDH (PDB ID 1U8F) (38) was visually inspected in PYMOL for suitable loops that could be altered for insertion of a FlAsH biarsenical-binding tetracysteine motif. Residue Gly58 of the loop immediately preceding the β4 strand was identified as a candidate and the FlAsH biarsenical-binding tetracysteine motif RWCCPGCCK was modeled into position using manual model building and geometry optimization in Coot (39). Initial models suggested that the insertion of RWCCPGCCK in place of Gly58 would place the FlAsH tag approximately 15 to 20 Å from the heme-binding His53. The FlAsH biarsenical-binding tetracysteine motif RWCCPGCCK was chosen due to report of the sequence as an improved version of the TC tag with optimized flanking residues as reported in (40). The nucleotide sequence coding for ‘RWCCPGCCK’ was ‘CGATGGTGCTGCCCGGGCTGCTGCAAG’ and was inserted into the coding sequence of human GAPDH. We used pRK5-HA vector to clone the TC-hGAPDH for the mammalian expression plasmid and pGST-parallel1 vector to clone the TC-hGAPDH for the bacterial expression plasmid. The H53A point mutation was made on both the mammalian and bacterial expression plasmids containing the TC-hGAPDH and used in our experiments.

### Growth of HEK293T cells, plasmid transfection and expression of human proteins

HEK293T cells (ATCC # CRL-3216) were cultured in tissue culture treated plates containing Dulbecco’s Modified Eagle’s Medium (DMEM) (ATCC # 30-2002) containing 10% fetal bovine serum (Gibco) until 60% confluent after which cells were transfected using Lipofectamine 2000 (Invitrogen # 11668019) with mammalian expression plasmids expressing pRK5-HA-TC-hGAPDH, pCMV3-hIDO1-FLAG (Sino Biologicals # HG11650-CF) as required. Protein expression was allowed for 48h in absence or presence of 1 mM δ-ALA and 100 µM Fe-cit until the plates reach full confluency. When needed cycloheximide (Chx) (Sigma # C7698) was used to treat cells at 5 μg/ml for 12 h to inhibit further protein synthesis. Cells were then utilized for experiments which in some cases involved treatment with succinyl acetone (SA) (Sigma # D1415) at 400 μM to inhibit further heme synthesis, removal of δ-ALA and Fe-cit heme precursors, and/or addition of NOC-18 NO donor at indicated doses and time points.

### Generation of ^14^C-labelled heme in cells

HEK293T cells were grown in DMEM/F-12 medium without Glycine (Caisson Labs # DFP04) with 10% heme depleted serum and 15 µM of ^14^C-δ-ALA and 10 µM Fe-cit for 48h post transfection of FLAG-IDO1 and HA-TC-hGAPDH plasmids. At this stage the proteins are expressed and have ^14^C-labeled heme incorporated into them. Chx was used to treat cells at 5 μg/ml for 12 h to inhibit further protein synthesis. Cells were lysed using 50 mM Tris-HCl pH 7.4 buffer with 0.1% Triton X-100, 5 mM Na-molybdate, and EDTA-free protease inhibitor cocktail (Roche). Protein concentration was measured using the Bradford method (Bio-Rad # 500-0006). Immunoprecipitation pull-downs were performed using 1 mg of whole cell extracts with anti-HA antibody (Invitrogen # 26183). Protein G agarose beads (Millipore # 16-201) were used to pull down the antibody– protein complex. The beads were washed well with lysis buffer, the 1.5-ml tubes were inserted into 5 ml scintillation vials and 4 ml of scintillation fluid (Liquiscint, National Diagnostics # LS-121) was added to each vial. The method for measuring ^14^C heme counts using a scintillation counter is described in (24).

### Treatment of cells with NOC-18

NOC-18 (Dojindo Molecular Technologies # N379-12) was freshly dissolved in cell culture grade sterile PBS at a stock concentration of 10 mM and added to phenol red– free cell culture medium containing 10% serum, 400 µM SA and L-Trp at 2 mM final concentration. This medium was used to treat the cells for the indicated time points. The pH of the medium was measured to be 7.0; the reported half-life of NOC-18 at pH 7 at 37 °C was 13 h (Dojindo Molecular Technologies Inc. website for NOC-18).

### Treatment of cells with radicicol

Hsp90 inhibitor radicicol (Sigma # R2146) at 10µM was used to pretreat cells for 6 h before treatment with NOC-18. After pretreatment, cells were maintained in the presence of 10 μM radicicol during NOC-18 treatment for the indicated time points and doses.

### IDO1 activity assay in cell supernatants

The activity of IDO1 in cell supernatants was measured as described in (41) with modifications. Briefly, 100 μg of cell supernatant protein was added to a 200 μl reaction containing 50 mM potassium phosphate buffer (pH 6.5), 20 mM ascorbate, 10 μM methylene blue, 100 μg/ml catalase, and 1 mM L-Trp and incubated at 37 ° C for 30 min. The reactions were stopped by adding 200 μl of 30% trichloroacetic acid to denature the proteins. The next steps were followed as per the previously described colorimetric detection method of Kyn using Ehrlich’s reagent. In some cases, the cell supernatants were given heme prior to assay. Supernatants (200 μg protein) were added to 6 μM heme, incubated at room temperature for 1 h, passed through a desalting column (Sephadex G-25 resin), and then assayed for activity as described above.

### Staining live cells expressing HA-TC-hGAPDH protein with FIAsH-EDT_2_ dye

HEK293T cells expressing HA-TC-hGAPDH protein with or without FLAG-IDO1 in black walled 96-well plate (Greiner Bio-One # 655090) were stained with FlAsH-EDT_2_. Briefly, the medium was aspirated out from the wells and washed once with cell culture PBS. FlAsH-EDT_2_ (Cayman chemicals # 20704) was dissolved in DMSO to a stock of 20 mM. The cells were stained with FlAsH-EDT_2_ diluted in opti-MEM medium at a final concentration of 5 μM at 37 °C for 30 min. The dye was aspirated out from the wells and washed twice with cell culture PBS. Phenol red free DMEM with or without NOC-18 and SA was added to the wells and the kinetic studies were performed.

### Live cell heme binding/transfer kinetics using FIAsH-EDT_2_ labeled HA-TC-hGAPDH

The heme binding kinetics onto HA-TC-hGAPDH or heme loss from HA-TC-hGAPDH under different experimental conditions was monitored in live cells stained with FIAsH-EDT_2_. We used FlexStation 3 (Molecular Devices) to perform the kinetic studies. The FIAsH-EDT_2_ was excited at 508 nm and emission was recorded at 528 nm. The relative fluorescence units (RFU) readings were recorded from the bottom of the plate, at 100 reads per well per reading at medium sensitivity for the PMT settings for the entire duration of the experiment and the temperature was maintained at 37 °C inside the instrument.

### Measurement of heme and total porphyrin from cell extract

Total heme and porphyrin from cell extracts were measured as described in (42). Briefly, 60 µl of cell supernatant was mixed with 240 µl of heme chromogen reagent (40:60 pyridine:H_2_O, 200 mM NaOH), and heme iron was reduced by adding a few grains of sodium dithionite. Heme chromogen formation was monitored at 556 nm, and its concentration was calculated using extinction co-efficient of 34.6 mM^-1^ cm^-1^. To measure total soluble porphyrin (soluble heme plus soluble porphyrin) in the cell supernatants, a fluorometric method was used (42). Briefly, in a micro-centrifuge tube, 980 µl of 2 M oxalic acid was mixed with 20 µl of cell supernatant and boiled for 1h. The tubes were allowed to cool to room temperature, after which porphyrin was measured by its fluorescence emission at 662 nm (excitation 400 nm) relative to standard curves generated with freshly prepared authentic heme solutions that had been subjected to similar oxalic acid treatment.

### Labile heme measurements in cells using a fluorescent heme sensor

The pcDNA3.1-eGFP-HS1 plasmid was a gift from Dr. Amit Reddi, Georgia Institute of Technology. Its composition and use were as described previously (8,24,26). Briefly, HEK293T cells cultured in DMEM with FBS (10%) were transfected with pcDNA3.1-eGFP-HS1 using Lipofectamine 2000. After 16 h the medium was replaced with medium containing δ-ALA (500 μM) + Fe-cit (200 μM). At various times, culture dishes were placed on ice, cells were removed, washed with ice cold PBS, re-suspended in PBS + glucose at 4 °C, and then taken to measurement. mKATE2 positive cells (1 × 10^6^ per condition) were analyzed for the intensity of their GFP and mKATE2 fluorescence on a BD LSR Fortessa instrument (BD Biosciences) (mKATE2 excitation 588 nm, emission 620 nm; GFP excitation 488 nm, emission 510 nm). Data were analyzed using FlowJo v10 (BD Biosciences) and were either graphed as a distribution of the cell number versus GFP:mKATE2 fluorescence ratio at each time point, or as the cell number with a fluorescence ratio of 0.5 versus time.

### Western blots

To check protein expressions across all samples in all experiments, we lysed cells using 50 mM Tris-HCl pH 7.4 buffer with 0.1% Triton X-100, 5 mM Na-molybdate, and EDTA-free protease inhibitor cocktail (Roche). Protein concentration was measured using the Bradford method (Bio-Rad # 500-0006). In each sample 10 μg of cell extract proteins was boiled in Laemmli buffer, resolved onto 10% or 15% SDS-PAGE, and transferred to polyvinylidene difluoride (PVDF) membrane (Bio-Rad # 1620177) and probed for proteins of interest. Western blot was performed with anti-IDO1 (Santa Cruz Biotechnology # sc-137012; dilution 1:1000), anti-HA (Santa Cruz Biotechnology # sc-7392; dilution 1:1000), and anti-α-Tubulin (Santa Cruz Biotechnology # sc-5286; dilution 1:1000). The proteins were detected using chemiluminescence using horseradish peroxidase–conjugated secondary antibodies of anti-mouse (Bio-Rad # 170-6516, dilution 1:10,000) origin and enhanced chemiluminescence substrate (Thermo Scientific # 32106). The images were acquired using a ChemiDoc MP System from Bio-Rad.

### NO release rate from NOC-18

The NO-mediated conversion of oxyhemoglobin to methemoglobin was used to determine the rate of NO release from NOC-18 at 37 °C. Various concentrations of NOC-18 were added to cuvettes that contained phenol red–free DMEM, 10% fetal bovine serum, 2 mM L-Trp, and 10 μM oxy-hemoglobin. The absorbance gain at 401 nm was recorded per minute over a 3 h period for each concentration of NOC-18. The rate of NO release was calculated using the difference extinction coefficient of 38 mM^−1^ cm^−1^ (17).

### Purification of hGAPDH and TC-hGAPDH proteins from bacteria

BL21DE3 *E. coli* cells that had been transformed with the hGAPDH-GST and TC-hGAPDH-GST expression plasmids were used to inoculate 500 ml cultures of terrific broth (2 liters total for each) containing 100 µg/ml carbenicilin as detailed in (24). The cultures were grown at 37 °C to an OD of 0.6-0.8 at 37 °C and then were cooled to RT before being induced with 1 mM IPTG. Cultures were shaken at RT and 250 rpm for 48 h before being harvested by centrifugation. Cells were pelleted then re-suspended in lysis buffer (50 mM Tris HCl pH 7.4, 10 mM MgCl_2_, 150 mM NaCl, 10% glycerol, 2 mM BME, protease inhibitor cocktail, DNase I, 1 mg/ml lysozyme) and flash frozen in liquid nitrogen. Lysates were then thawed overnight at 4 °C with rotation before being sonicated and spun in a centrifuge at 4 °C, with the supernatant being collected. Glutathione agarose beads (MCLAB # GAB-300) were equilibrated with wash buffer (50 mM Tris HCl pH 8, 10 mM MgCl_2_, 150 mM NaCl, 5% glycerol, 2 mM BME) and the supernatant was mixed with the beads by gentle shaking for 2-3 h at 4 °C (25 ml beads per liter equivalent of cells). The beads were then washed until the flow through is clear. Thrombin was added to the beads, gently mixed, and then allowed to incubate overnight at 4 °C. After the thrombin incubation the GAPDH protein was eluted from beads using cold wash buffer. Benzamidine sepharose beads were then incubated with the eluted protein for 30 min to remove thrombin. The protein solution was then concentrated in a 10 kDa cut-off centrifugal filter, buffer exchanged if desired, and stored at -80 °C. Purity was assessed by SDS-PAGE and the protein concentration determined by Bradford assay (Bio-Rad).

### Heme titration of purified TC-hGAPDH in plate reader

TC-hGAPDH was concentrated to 50-100 µM (monomer) and dialyzed into labeling buffer (50 mM MOPS, 150 mM NaCl, pH=7.4). The protein was reduced using 2 mM TCEP for 1 h on ice, then FIAsH-EDT_2_ (dissolved in DMSO) was added in a 2:1 ratio with GAPDH monomer. Protein was incubated for either 30 min at 37 °C or overnight at 4 °C protected from light. To remove excess dye, the sample was passed through a desalting column (PD spin trap G-25) equilibrated with labeling buffer. Heme solutions were prepared by dissolving hemin chloride into DMSO and then passing through a 0.22 µM nylon filter. The concentration of heme was determined by measuring absorbance at 404 nm (170,000 M^-1^ cm^-1^). TC-hGAPDH (1 µM or 20 nM) was added to each well and varying doses of heme were added. The sample was mixed briefly on an orbital shaker (400 rpm, 10 sec) then allowed to equilibrate in the dark for 30 seconds. An endpoint reading of the fluorescent signal was then measured (excitation 508 nm, emission 528 nm).

### Heme titration of purified TC-GAPDH in a conventional fluorimeter

TC-hGAPDH (1 µM or 5 µM) protein was added to a fluorescence cuvette and an emission scan was recorded (excitation 508 nm) on a Hitachi F-2500 fluorescence spectrophotometer. Heme in different concentrations were then added to the protein and incubated for 30 s protected from light before the emission scan was measured at 528 nm.

### Heme titration of TC-GAPDH in cell supernatants

HEK293T cells were grown in DMEM containing 10% serum. Cells were allowed to express the HA-TC-hGAPDH for 48 h post transfection of plasmid. Cells were then incubated with FlAsH in Opti-MEM for 30 min at RT protected from light. Afterwards, cells were washed three times with phenol red-free DMEM. Cells were lysed using ice cold 50 mM Tris-HCl pH 7.4 buffer with 0.1% Triton X-100, 5 mM Na-molybdate, and EDTA-free protease inhibitor cocktail (Roche) and lysates spun at 4 °C at 10,000 x g for 10 min to yield supernatants for analysis. Protein concentration was measured using the Bradford method (Bio-Rad # 500-0006). 0.08 mg/mL of supernatant protein was mixed in buffer (50 mM MOPS, 150 mM NaCl, pH 7.4) at RT with varying concentrations of heme (0-2 µM) which were prepared from a stock solution of hemin chloride in DMSO. After mixing the reaction was allowed to equilibrate for 30 min in the dark. The end point fluorescence signals were measured in the 96-well plate reader at RT using excitation at 508 nm and emission at 528 nm.

### Statistical analyses

All experiments were done in three independent trials, with three replicates per trial. The results are presented as the mean of the three trial values ± standard deviation. The statistical test used to measure significance (p-values) was student’s t-test in the software GraphPad Prism (v9).

## Data Availability

All data are contained in this manuscript or are available from the authors upon request.

## Supporting information

This article contains supporting information.

## Funding and additional information

This work was supported by National Institutes of Health grants R01 GM130624 and R01 GM148664 (to D.J. Stuehr) and R35 GM128595 (to R.C. Page). The content is solely the responsibility of the authors and does not necessarily represent the official views of the National Institutes of Health.

## Acknowledgments

We thank members of the Stuehr lab for helpful discussion.

## Author Contributions

Pranjal Biswas, Joseph Palazzo, Simon Schlanger, Dhanya Thamaraparambil Jayaram, Sidra Islam, Richard C. Page: Methodology, Writing - Review & Editing, Funding acquisition, Validation, Formal Analysis, Investigation. Dennis J. Stuehr: Conceptualization, Methodology, Validation, Formal Analysis, Investigation, Data Curation, Writing - Original Draft, Writing - Review & Editing, Visualization, Supervision, Project administration, Funding acquisition.

## References

1. Harvey, J. W., and Beutler, E. (1982) Binding of heme by glutathione S-transferase: a possible role of the erythrocyte enzyme. Blood 60, 1227–1230

2. Iwahara, S., Satoh, H., Song, D. X., Webb, J., Burlingame, A. L., Nagae, Y., and Muller-Eberhard, U. (1995) Purification, characterization, and cloning of a heme-binding protein (23 kDa) in rat liver cytosol. Biochemistry 34, 13398–13406

3. Taketani, S., Adachi, Y., Kohno, H., Ikehara, S., Tokunaga, R., and Ishii, T. (1998) Molecular characterization of a newly identified heme-binding protein induced during differentiation of urine erythroleukemia cells. J Biol Chem 273, 31388–31394

4. Vincent, S. H., and Muller-Eberhard, U. (1985) A protein of the Z class of liver cytosolic proteins in the rat that preferentially binds heme. J Biol Chem 260, 14521–14528

5. Donegan, R. K., Moore, C. M., Hanna, D. A., and Reddi, A. R. (2019) Handling heme: The mechanisms underlying the movement of heme within and between cells. Free Radic Biol Med 133, 88–100

6. Tupta, B., Stuehr, E., Sumi, M. P., Sweeny, E. A., Smith, B., Stuehr, D. J., and Ghosh, A. (2022) GAPDH is involved in the heme-maturation of myoglobin and hemoglobin. FASEB J 36, e22099

7. Biswas, P., Dai, Y., and Stuehr, D. J. (2022) Indoleamine dioxygenase and tryptophan dioxygenase activities are regulated through GAPDH- and Hsp90-dependent control of their heme levels. Free Radic Biol Med 180, 179–190

8. Dai, Y., Sweeny, E. A., Schlanger, S., Ghosh, A., and Stuehr, D. J. (2020) GAPDH delivers heme to soluble guanylyl cyclase. J Biol Chem 295, 8145–8154

9. Chakravarti, R., Aulak, K. S., Fox, P. L., and Stuehr, D. J. (2010) GAPDH regulates cellular heme insertion into inducible nitric oxide synthase. Proc Natl Acad Sci U S A 107, 18004–18009

10. Dai, Y., Fleischhacker, A. S., Liu, L., Fayad, S., Gunawan, A. L., Stuehr, D. J., and Ragsdale, S. W. (2022) Heme delivery to heme oxygenase-2 involves glyceraldehyde-3-phosphate dehydrogenase. Biol Chem 403, 1043–1053

11. Morishima, Y., Lau, M., Pratt, W. B., and Osawa, Y. (2023) Dynamic cycling with a unique Hsp90/Hsp70-dependent chaperone machinery and GAPDH is needed for heme insertion and activation of neuronal NO synthase. J Biol Chem 299, 102856

12. Fleischhacker, A. S., and Ragsdale, S. W. (2018) An unlikely heme chaperone confirmed at last. J Biol Chem 293, 14569–14570

13. Stuehr, D. J., Dai, Y., Biswas, P., Sweeny, E. A., and Ghosh, A. (2022) New roles for GAPDH, Hsp90, and NO in regulating heme allocation and hemeprotein function in mammals. Biol Chem 403, 1005-1015

14. Medlock, A. E., and Dailey, H. A. (2022) New Avenues of Heme Synthesis Regulation. Int J Mol Sci 23

15. Stuehr, D. J., Biswas, P., Dai, Y., Ghosh, A., Islam, S., and Jayaram, D. T. (2023) A natural heme deficiency exists in biology that allows nitric oxide to control heme protein functions by regulating cellular heme distribution. Bioessays 45, e2300055

16. Lundberg, J. O. (2023) Nitric oxide achieves master regulator status. Bioessays 45, e2300089

17. Biswas, P., and Stuehr, D. J. (2023) Indoleamine dioxygenase and tryptophan dioxygenase activities are regulated through control of cell heme allocation by nitric oxide. J Biol Chem 299, 104753

18. Waheed, S. M., Ghosh, A., Chakravarti, R., Biswas, A., Haque, M. M., Panda, K., and Stuehr, D. J. (2010) Nitric oxide blocks cellular heme insertion into a broad range of heme proteins. Free Radic Biol Med 48, 1548–1558

19. Dai, Y., Faul, E. M., Ghosh, A., and Stuehr, D. J. (2022) NO rapidly mobilizes cellular heme to trigger assembly of its own receptor. Proc Natl Acad Sci U S A 119

20. Sumi, M. P., Tupta, B., and Ghosh, A. (2022) Nitric Oxide Trickle Drives Heme into Hemoglobin and Muscle Myoglobin. Cells 11

21. Ghosh, A., Sumi, M. P., Tupta, B., Okamoto, T., Aulak, K., Tsutsui, M., Shimokawa, H., Erzurum, S. C., and Stuehr, D. J. (2022) Low levels of nitric oxide promotes heme maturation into several hemeproteins and is also therapeutic. Redox Biol 56, 102478

22. Adams, S. R., Campbell, R. E., Gross, L. A., Martin, B. R., Walkup, G. K., Yao, Y., Llopis, J., and Tsien, R. Y. (2002) New biarsenical ligands and tetracysteine motifs for protein labeling in vitro and in vivo: synthesis and biological applications. J Am Chem Soc 124, 6063–6076

23. Ito, H., Nishio, Y., Hara, T., Sugihara, H., Tanaka, T., and Li, X. K. (2018) Oral administration of 5-aminolevulinic acid induces heme oxygenase-1 expression in peripheral blood mononuclear cells of healthy human subjects in combination with ferrous iron. Eur J Pharmacol 833, 25–33

24. Sweeny, E. A., Singh, A. B., Chakravarti, R., Martinez-Guzman, O., Saini, A., Haque, M. M., Garee, G., Dans, P. D., Hannibal, L., Reddi, A. R., and Stuehr, D. J. (2018) Glyceraldehyde-3-phosphate dehydrogenase is a chaperone that allocates labile heme in cells. J Biol Chem 293, 14557–14568

25. Hoffmann, L. S., Schmidt, P. M., Keim, Y., Hoffmann, C., Schmidt, H. H., and Stasch, J. P. (2011) Fluorescence dequenching makes haem-free soluble guanylate cyclase detectable in living cells. PLoS One 6, e23596

26. Hanna, D. A., Harvey, R. M., Martinez-Guzman, O., Yuan, X., Chandrasekharan, B., Raju, G., Outten, F. W., Hamza, I., and Reddi, A. R. (2016) Heme dynamics and trafficking factors revealed by genetically encoded fluorescent heme sensors. Proc Natl Acad Sci U S A 113, 7539–7544

27. Sassa, S., and Kappas, A. (1983) Hereditary tyrosinemia and the heme biosynthetic pathway. Profound inhibition of delta-aminolevulinic acid dehydratase activity by succinylacetone. J Clin Invest 71, 625–634

28. Tschudy, D. P., Hess, R. A., and Frykholm, B. C. (1981) Inhibition of delta-aminolevulinic acid dehydrase by 4,6-dioxoheptanoic acid. J Biol Chem 256, 9915–9923

29. Seidler, N. W. (2013) Basic biology of GAPDH. Adv Exp Med Biol 985, 1–36

30. Barber, R. D., Harmer, D. W., Coleman, R. A., and Clark, B. J. (2005) GAPDH as a housekeeping gene: analysis of GAPDH mRNA expression in a panel of 72 human tissues. Physiol Genomics 21, 389–395

31. Tristan, C., Shahani, N., Sedlak, T. W., and Sawa, A. (2011) The diverse functions of GAPDH: views from different subcellular compartments. Cell Signal 23, 317–323

32. Chen, J. J. (2007) Regulation of protein synthesis by the heme-regulated eIF2alpha kinase: relevance to anemias. Blood 109, 2693–2699

33. Doty, R. T., Phelps, S. R., Shadle, C., Sanchez-Bonilla, M., Keel, S. B., and Abkowitz, J. L. (2015) Coordinate expression of heme and globin is essential for effective erythropoiesis. J Clin Invest 125, 4681–4691

34. Ignarro, L. J., Degnan, J. N., Baricos, W. H., Kadowitz, P. J., and Wolin, M. S. (1982) Activation of purified guanylate cyclase by nitric oxide requires heme. Comparison of heme-deficient, heme-reconstituted and heme-containing forms of soluble enzyme from bovine lung. Biochim Biophys Acta 718, 49–59

35. Ignarro, L. J., Adams, J. B., Horwitz, P. M., and Wood, K. S. (1986) Activation of soluble guanylate cyclase by NO-hemoproteins involves NO-heme exchange. Comparison of heme-containing and heme-deficient enzyme forms. J Biol Chem 261, 4997–5002

36. Hall, C. N., and Garthwaite, J. (2009) What is the real physiological NO concentration in vivo? Nitric Oxide 21, 92–103

37. Somasundaram, V., Basudhar, D., Bharadwaj, G., No, J. H., Ridnour, L. A., Cheng, R. Y. S., Fujita, M., Thomas, D. D., Anderson, S. K., McVicar, D. W., and Wink, D. A. (2019) Molecular Mechanisms of Nitric Oxide in Cancer Progression, Signal Transduction, and Metabolism. Antioxid Redox Signal 30, 1124–1143

38. Jenkins, J. L., and Tanner, J. J. (2006) High-resolution structure of human D-glyceraldehyde-3-phosphate dehydrogenase. Acta Crystallogr D Biol Crystallogr 62, 290–301

39. Emsley, P., Lohkamp, B., Scott, W. G., and Cowtan, K. (2010) Features and development of Coot. Acta Crystallogr D Biol Crystallogr 66, 486–501

40. Martin, B. R., Giepmans, B. N., Adams, S. R., and Tsien, R. Y. (2005) Mammalian cell-based optimization of the biarsenical-binding tetracysteine motif for improved fluorescence and affinity. Nat Biotechnol 23, 1308–1314

41. Takikawa, O., Kuroiwa, T., Yamazaki, F., and Kido, R. (1988) Mechanism of interferon-gamma action. Characterization of indoleamine 2,3-dioxygenase in cultured human cells induced by interferon-gamma and evaluation of the enzyme-mediated tryptophan degradation in its anticellular activity. J Biol Chem 263, 2041-2048

42. Albakri, Q. A., and Stuehr, D. J. (1996) Intracellular assembly of inducible NO synthase is limited by nitric oxide-mediated changes in heme insertion and availability. J Biol Chem 271, 5414–5421

