## Supporting Information for "Visualizing Mitochondrial Heme Flow through GAPDH to Targets in Living Cells and its Regulation by NO"


| Contents | Page number |
|---|---|
| Fig S1 | S-1 |
| Fig S2 | S-2 |
| Fig S3 | S-3 |
| Fig S4 | S-4 |
| Fig S5 | S-5 |
| Fig S6 | S-6 |
| Fig S7 | S-7 |
| Fig S8 | S-8 |
| Fig S9 | S-9 |
| Fig S10 | S-10 |

Fig S1. Amino acid sequence of human GAPDH with TC insert in red.

>TC-hGAPDH_PROTEIN sequence

MGKVKVGVNGFGRIGRLVTRAAFNSGKVDIVAINDPFIDLNYMVYMFQYDSTHGKFHRW
CCPGCCKTVKAENGKLVINGNPITIFQERDPSKIKWGDAGAEYVVESTGVFTTMEKAGAHL
QGGAKRVIISAPSADAPMFVMGVNHEKYDNSLKIISNASCTTNCLAPLAKVIHDNFGIVE
GLMTTVHAITATQKTVDGPSGKLWRDGRGALQNIIPASTGAAKAVGKVIPELNGKLTGMA
FRVPTANVSVVDLTCRLEKPAKYDDIKKVVKQASEGPLKGILGYTEHQVVSSDFNSDTHS
STFDAGAGIALNDHFVKLISWYDNEFGYSNRVVDLMAHMASKE

Fig S2. Representative heme titration of FlAsH-labeled TC-GAPDH in cell supernatant.

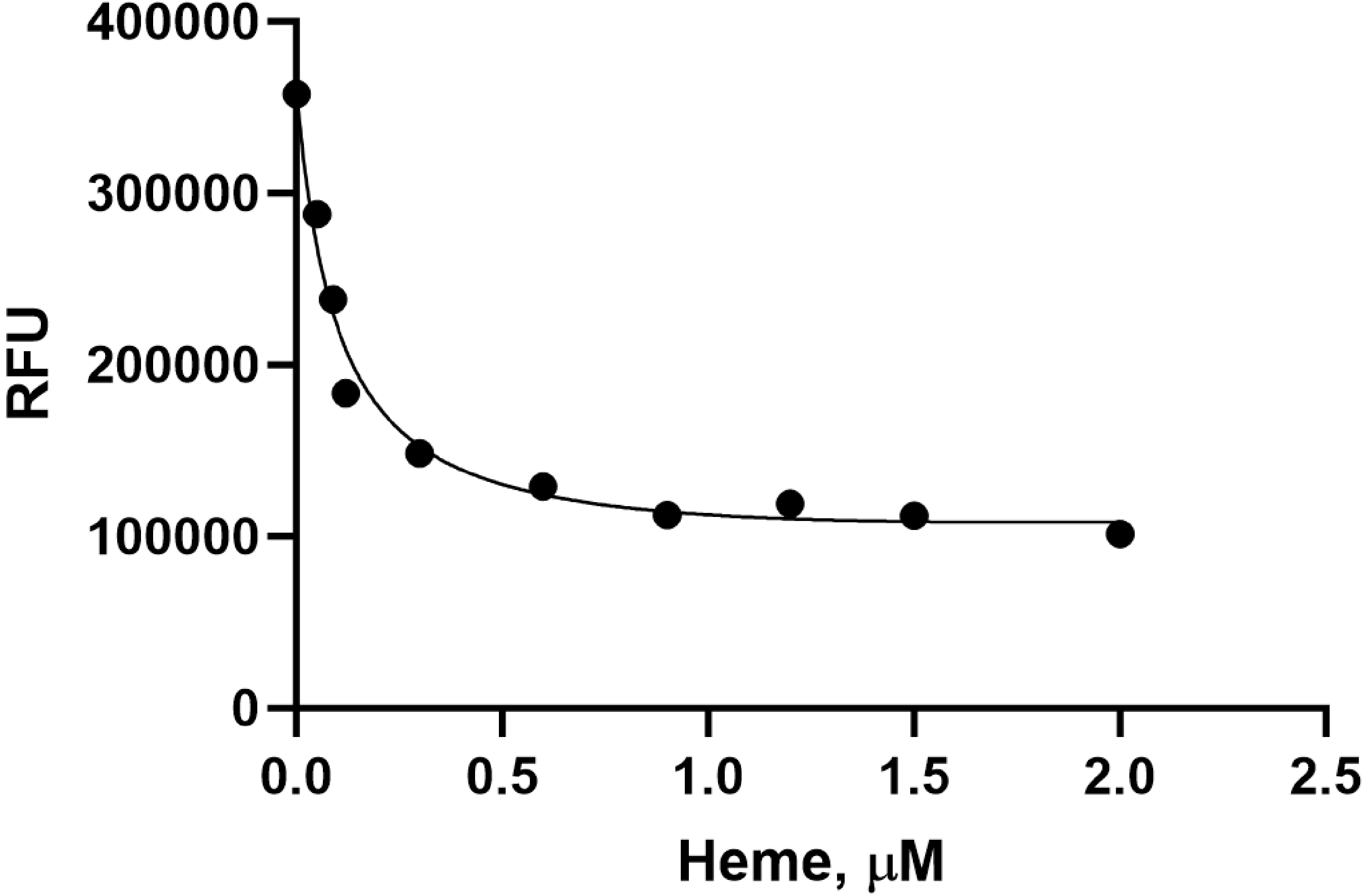

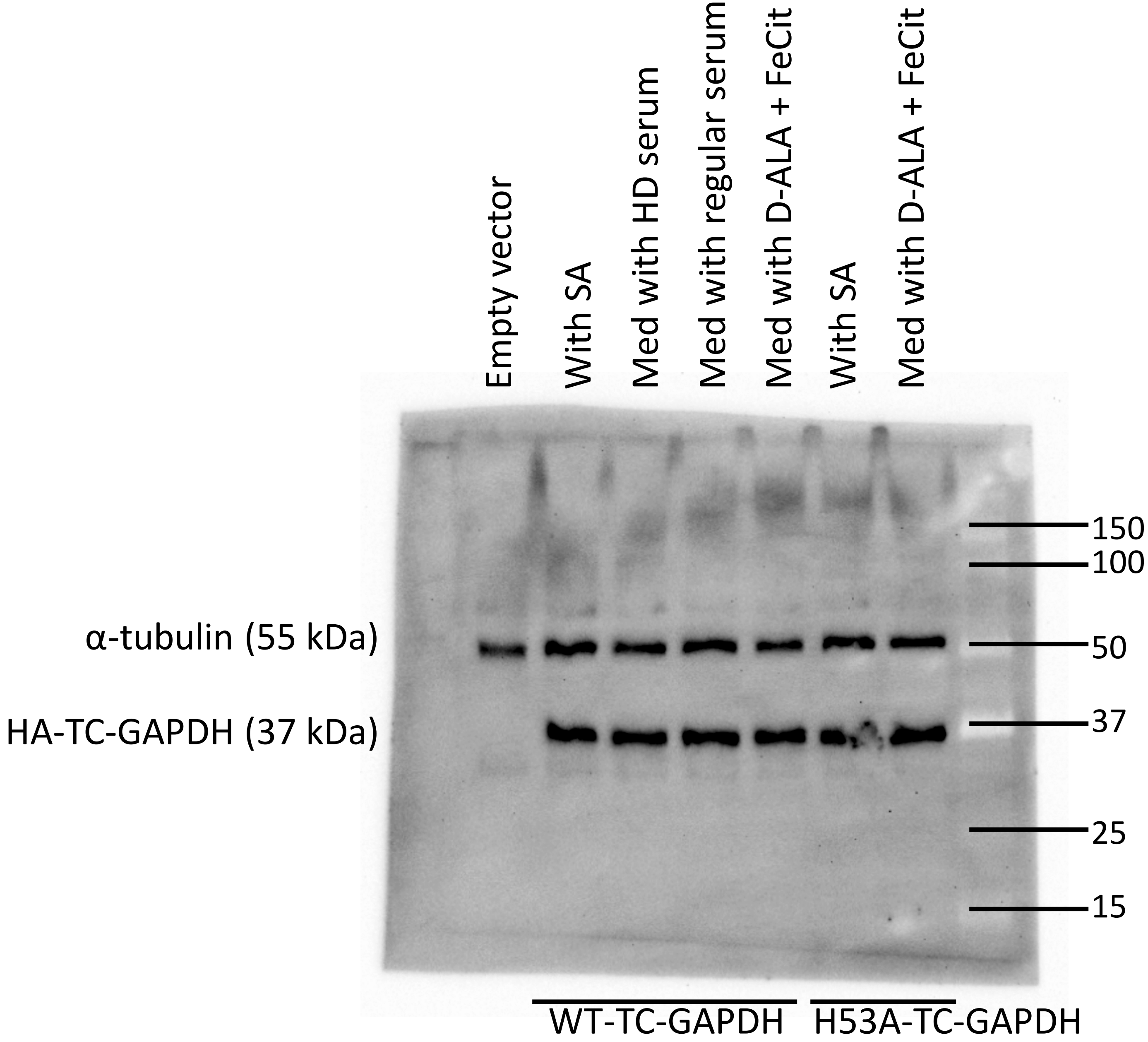


Fig S3. Representative WB for Fig 2.

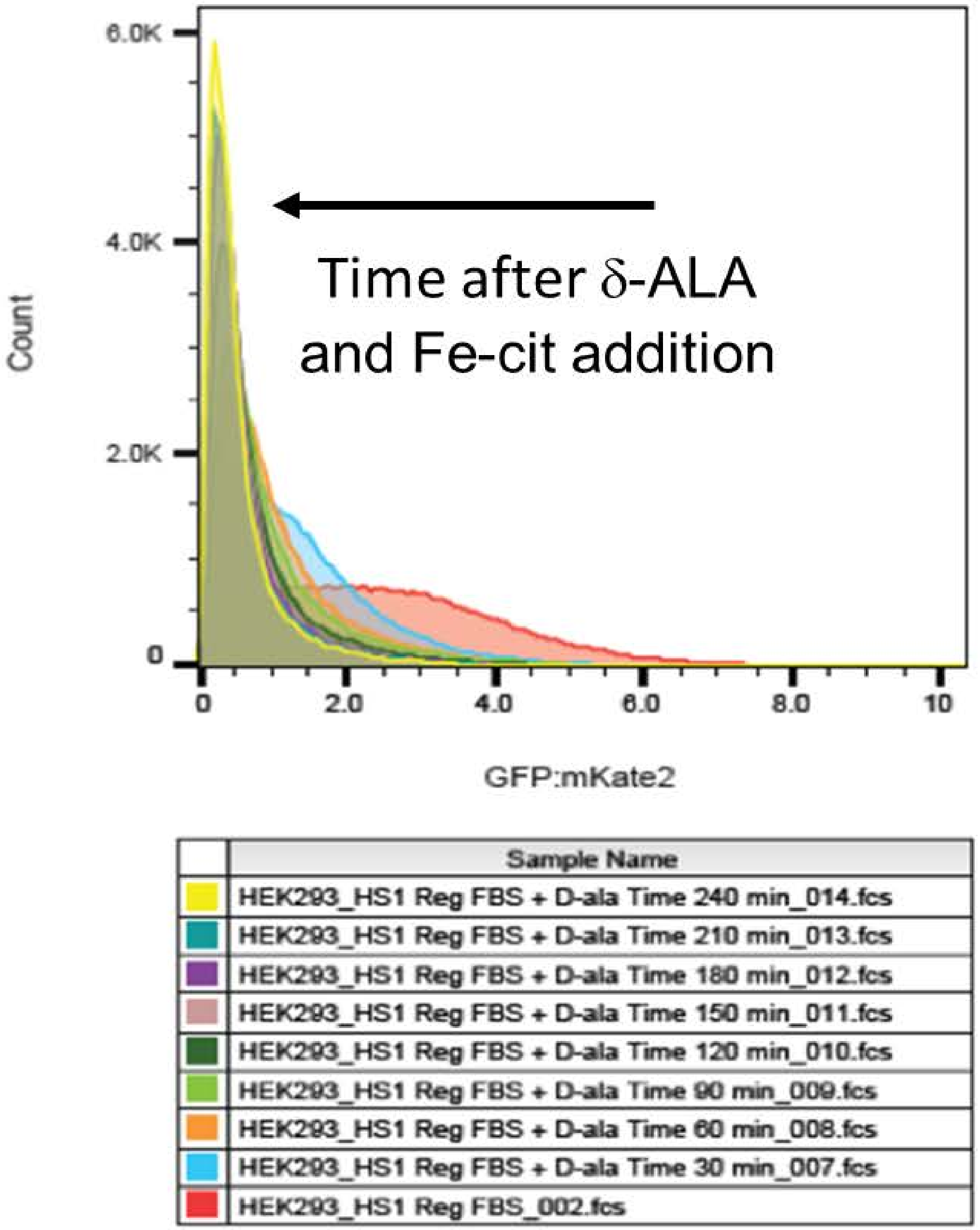


Fig S4. Change in cell number distribution according to their HS1 GFP:mKATE fluorescence ratios versus time after D-ALA and Fe-cit addition.

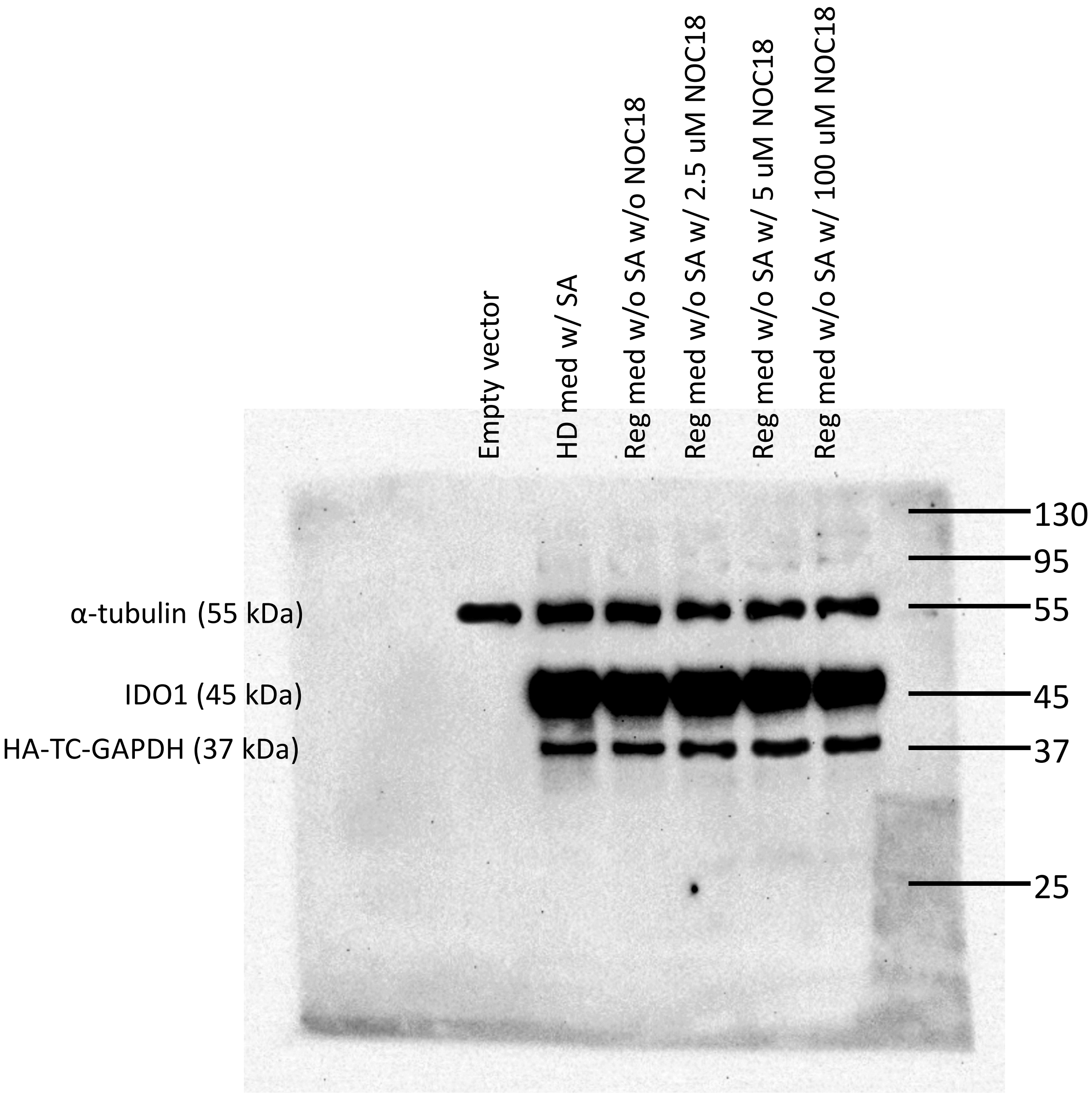


Fig S5. Representative WB for Fig 3.

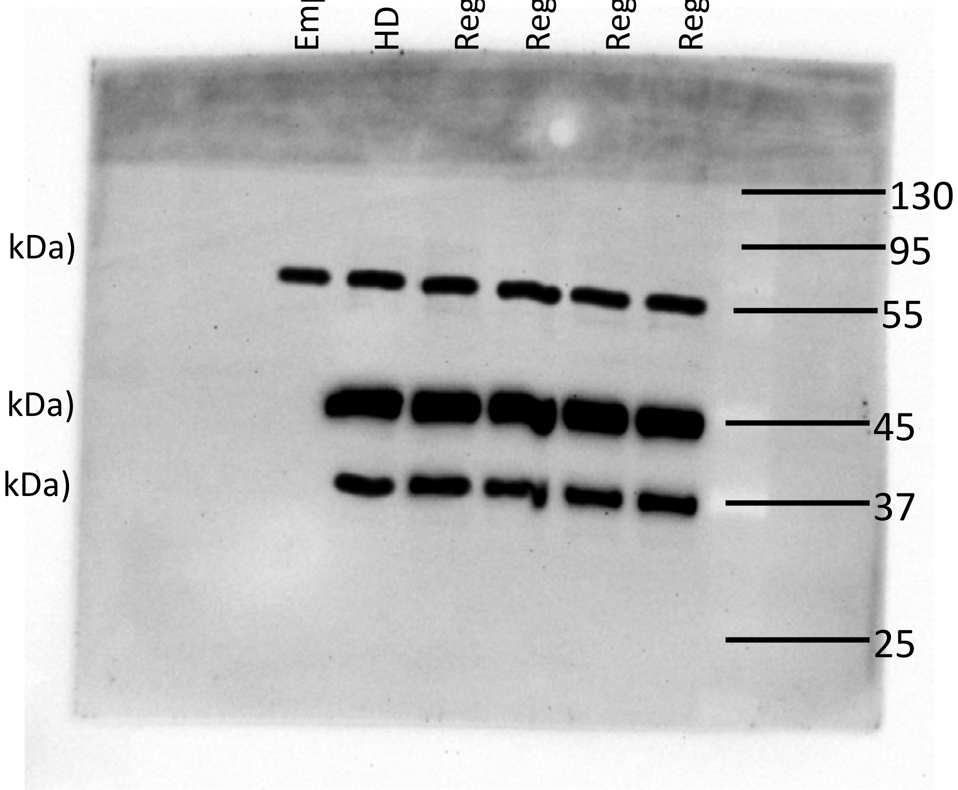


Fig S6. Representative WB for Fig 4.

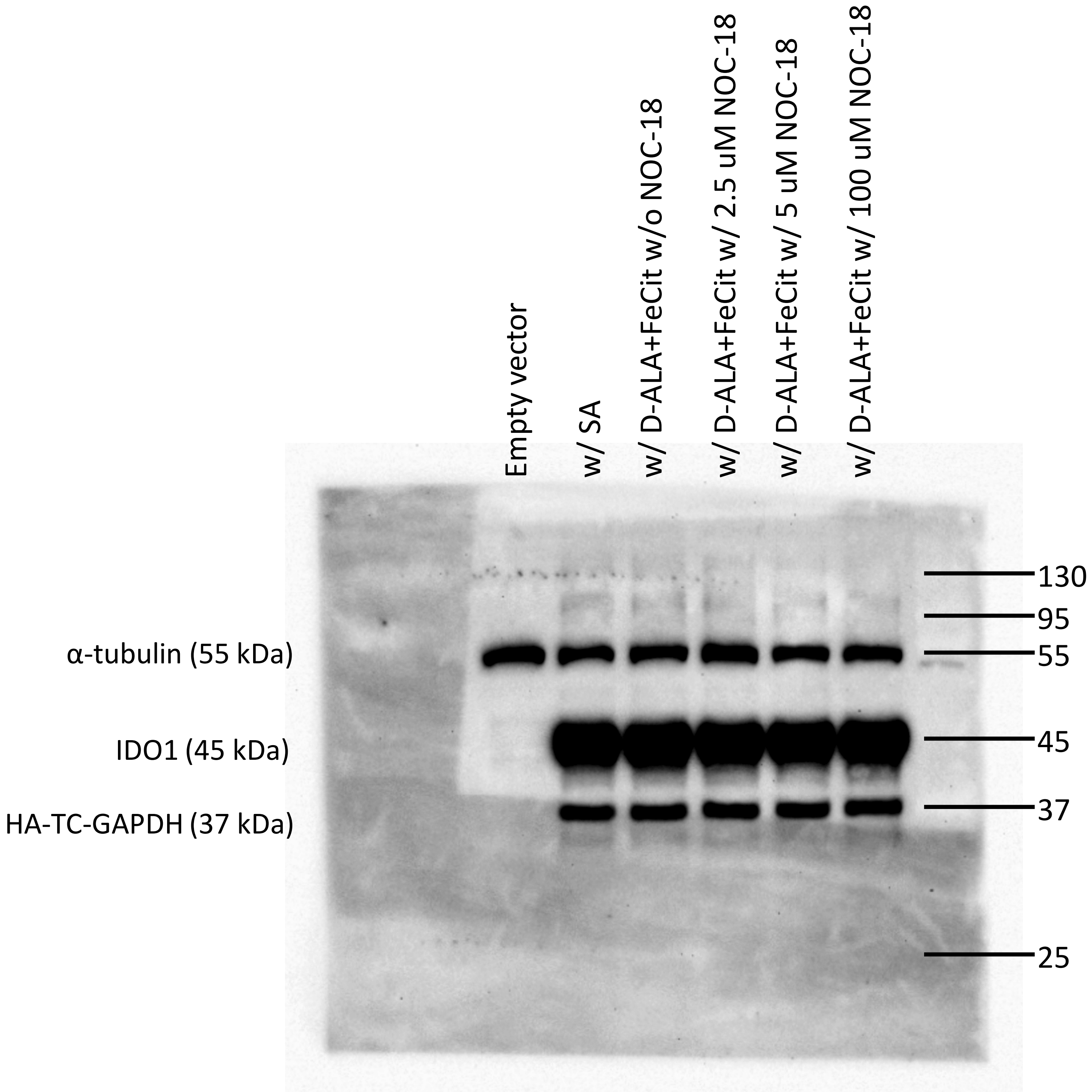


Fig S7. Representative WB for Fig 5.

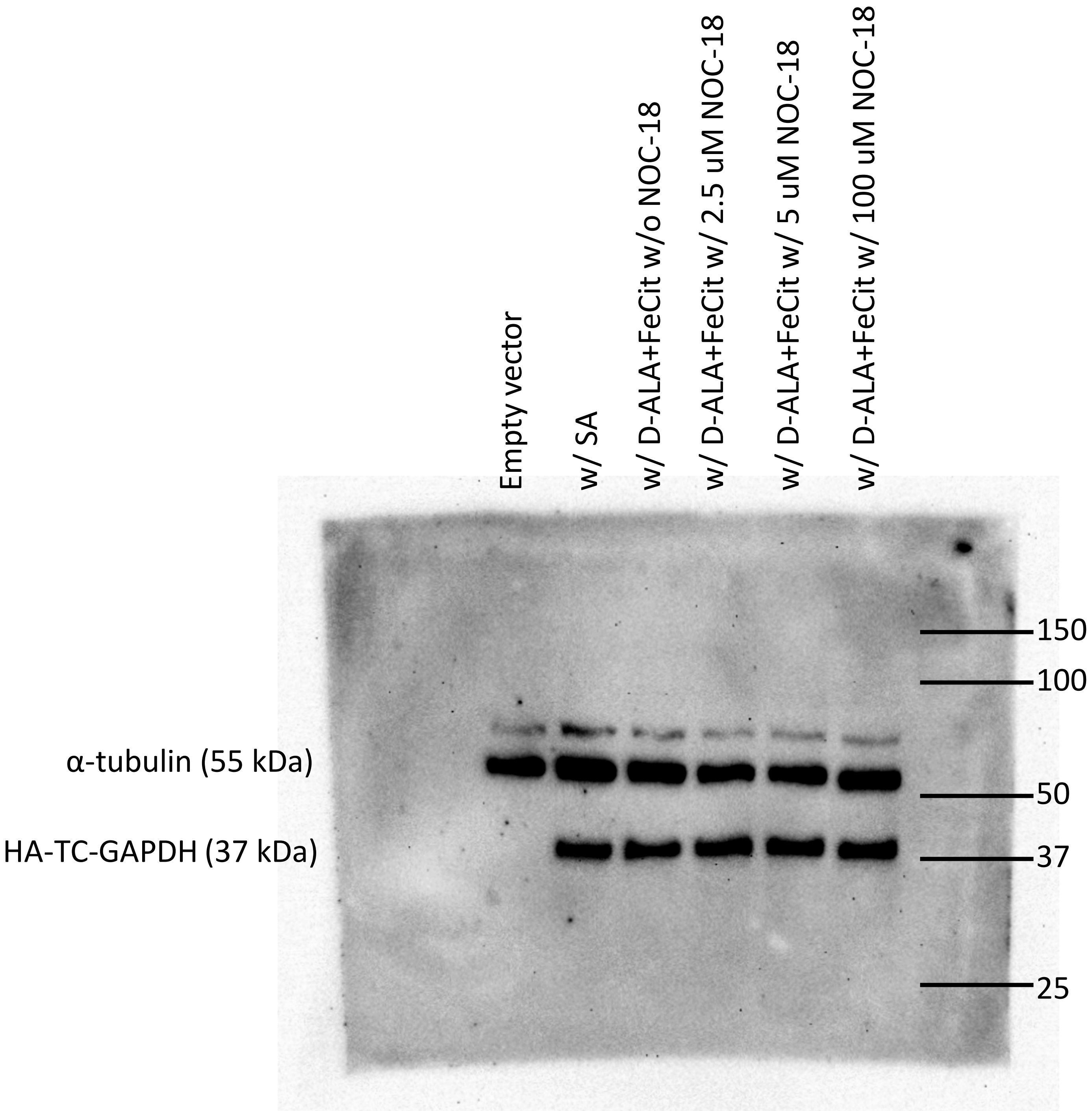


Fig S8. Representative WB for Fig 6.

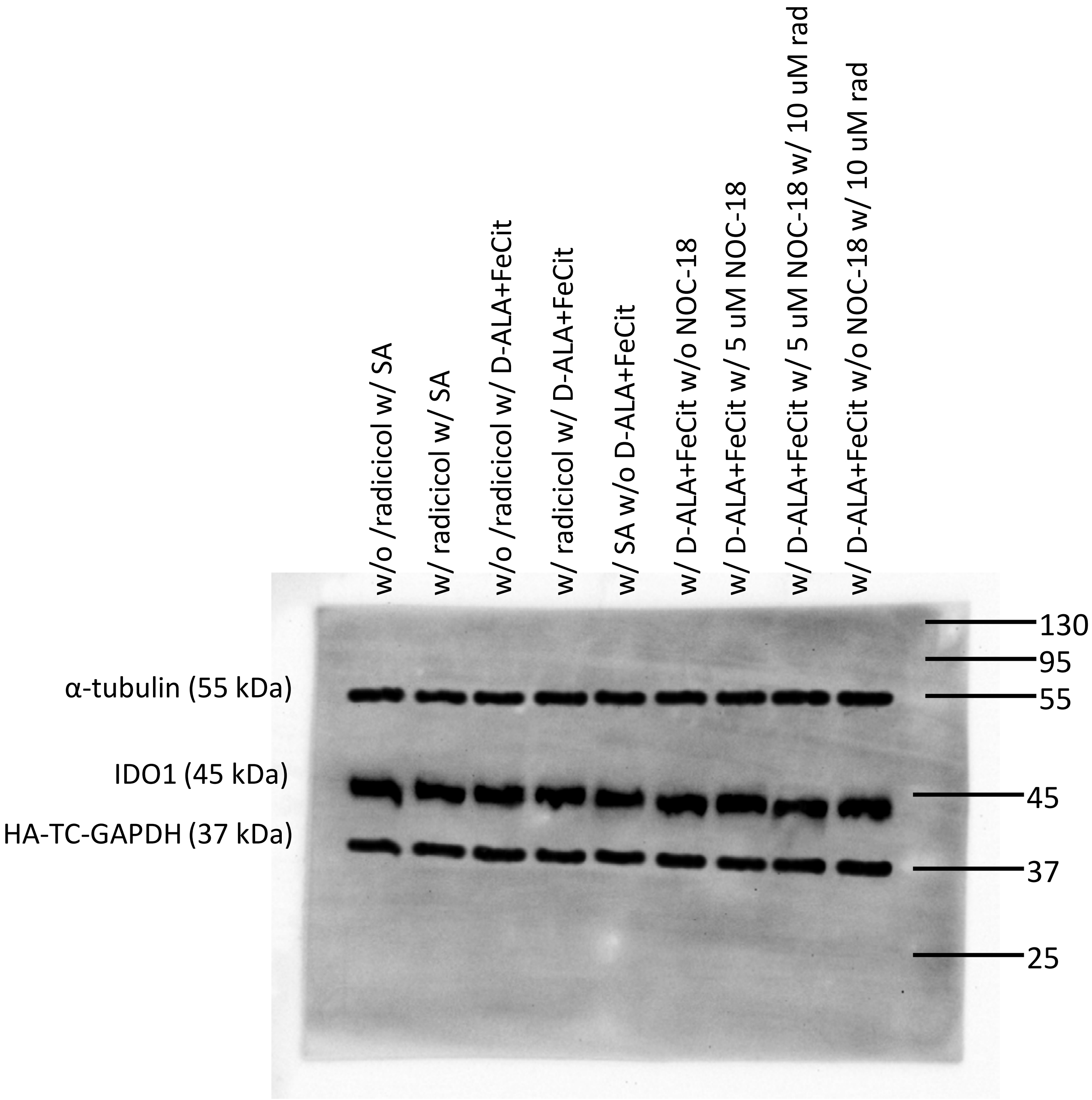


Fig S9. Representative WB for Fig 7.

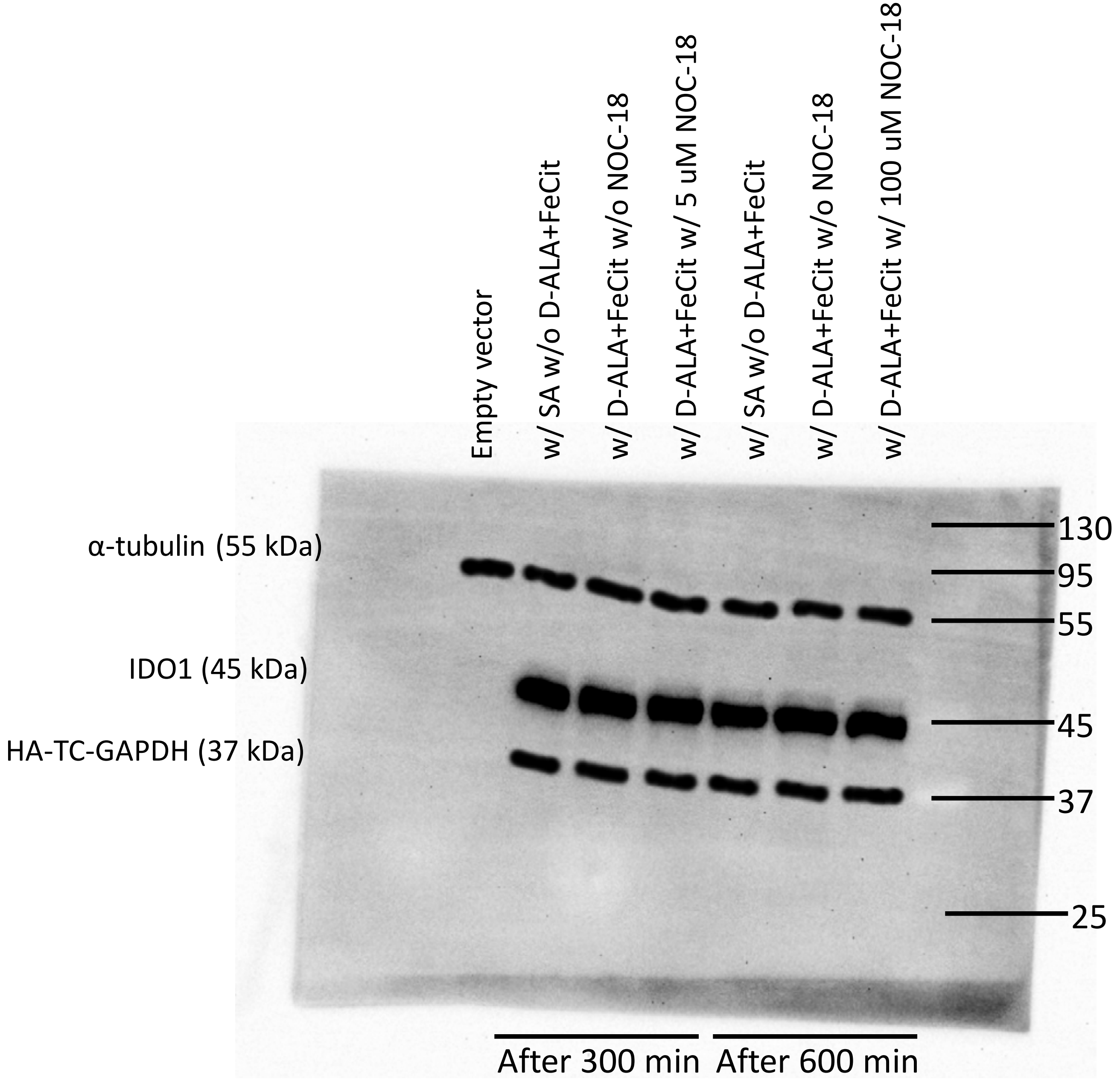


Fig S10. Representative WB for Fig 8.
